# Structural insights into HIV-2 CA lattice formation and FG-pocket binding revealed by single particle cryo-EM

**DOI:** 10.1101/2024.10.09.617312

**Authors:** Matthew Cook, Christian Freniere, Chunxiang Wu, Faith Lozano, Yong Xiong

## Abstract

One of the most striking features of HIV is the capsid; a fullerene cone comprised of the pleomorphic capsid protein (CA) which shields the viral genome from cellular defense mechanisms and recruits cellular cofactors to the virus. Despite significant advances in understanding the mechanisms of HIV-1 CA assembly and host factor interaction, HIV-2 CA remains poorly understood. By templating the assembly of HIV-2 CA on functionalized liposomes, we were able to determine high resolution structures of the HIV-2 CA lattice, including both CA hexamers and pentamers, alone and in complexes with peptides of host phenylalanine-glycine (FG)-motif proteins Nup153 and CPSF6. While the overall fold and mode of binding the FG-peptides are conserved with HIV-1, this study reveals distinctive structural features that define the HIV-2 CA lattice, potential differences in interactions with other host factors such as CypA, and divergence in the mechanism of formation of hexameric and pentameric CA assemblies. This study extends our understanding of HIV capsids and highlights an approach with significant potential to facilitate the study of lentiviral capsid biology.

## Introduction

Human immunodeficiency virus type 2 (HIV-2) was first identified in West Africa in 1986 among patients exhibiting AIDS symptoms. ^1,2^ The virus arose from a separate zoonotic spillover event than HIV-1, specifically deriving from a simian immunodeficiency virus infecting sooty mangabeys (SIV_smm_).^3,4^ While capable of progressing to AIDS when untreated, HIV-2 features slower average disease progression and lower viral loads compared to HIV-1.^5-7^

During HIV-1 or HIV-2 infection, the capsid is a necessary component which protects the viral genome and recruits host factors while trafficking to the nucleus.^8-10^ Acting as the shell of the virus core, the capsid is a major nexus for interactions with a large array of both pro-viral and restrictive host factors.^11,12^The vital role of the capsid and the many proteins targeting it apply significant selective pressure on the capsid protein (CA), and it is therefore broadly genetically fragile.^13,14^ Consequently, CA sequences are relatively well-conserved among lentiviruses.^15^On the other hand, host recognition of HIV-1 versus HIV-2 capsids is sometimes distinct, with notable implications on infectivity. ^16-18^Among many host factors studied, human TRIM5α ^18-25^and NONO^26^ are particularly noteworthy for their significant restriction of HIV-2, but limited effect on HIV-1.

The capsid is comprised of approximately ∼1,600 copies of the CA protein, which oligomerize into ∼250 hexamers and 12 pentamers (capsomers) to form the distinct fullerene cone.^11,27-30^ As the capsid is indispensable for successful infection, its pleomorphism is a key feature of HIV biology.^28,31-39^Understanding of the mechanisms of the accommodation of HIV-1 CA hexamers and pentamers in the capsid has recently advanced considerably alongside the ability to resolve structures of HIV-1 capsomeres in capsid-like lattices. It has been reported that the _58_TVGG_61_ loop of HIV-1 CA acts as a molecular “switch” between hexamer and pentamer, promoting remodeling of the hydrophobic core, gating of a host factor binding pocket, and adjusted inter-chain contacts among CA molecules.^38,39^

While crystal structures of the individual N-terminal domain (NTD)^40^and C-terminal domain (CTD)^41^of HIV-2 CA have been solved, the current methods are inadequate for probing the atomic-level details of HIV-2 CA lattice assembly and its interactions with host factors, as the formation of the mature HIV-2 CA lattice has proven to be more challenging compared to that of HIV-1 CA.^42^ As such, the structural details underscoring the biologic role of the assembled HIV-2 capsid lattice remain poorly characterized. Given the recent progress in developing approaches to understand HIV-1 CA,^43,44^HIV-2 CA affords an opportunity to illuminate broader lentiviral capsid biology and potentially provide future directions for pharmacological intervention.

In this study, we leveraged the approach of templating of HIV CA lattice assembly on liposomes,^44^together with stabilization by the small molecule host factor inositol hexakisphosphate (IP6),^44-48^to produce regular HIV-2 CA lattices. We report the first high-resolution structures of HIV-2 CA capsomers in their native mature lattice using single particle cryo-electron microscopy (cryo-EM). While the structures are broadly conserved, the mechanism of hexamer/pentamer switching appears different between HIV-2 and HIV-1, suggesting divergent evolution of these lentiviral capsid systems. In addition, we highlight that this system can be used to study host factor-CA lattice interactions via structural resolution of complexes of HIV-2 CA assemblies with peptides of phenylalanine-glycine (FG)-motif proteins. Altogether, we provide new insights into mechanisms of HIV-2 capsid assembly and host factor targeting.

## Results

### A capsid-like lattice of HIV-2 CA can be assembled via liposome templating

Understanding of HIV-1 capsid biology has expanded significantly alongside an array of tools and approaches for reproducing the mature CA lattice with *in vitro* assemblies.^27,28,43-45,49-52^While functionally and structurally conserved, HIV-2 CA is distinct, with differing effects on infectivity and host factor interaction.^5,16-26^As no effective approaches for determining atomic-level structures of the HIV-2 capsid exist, we sought to identify a platform for the assembly of HIV-2 CA into capsid-like particles (CLPs) and resolve high-resolution structures of the mature CA lattice via cryo-EM. Recent work demonstrated efficient assembly of HIV-1 CLPs via protein templating on liposomes decorated with Ni-NTA headgroups.^44^ We adopted this approach for HIV-2 CA assemblies.

Templating is achieved by association of His-tagged CA protein with the nickel-nitrilotriacetic acid (Ni-NTA) modified lipid head groups in the liposome; we therefore introduced a C-terminal His tag to HIV-2 GL-AN^53^CA with an intervening Gly-Ser-Ser linker. As such, the numbering used in this work is based on the HIV-2 GL-AN sequence, which is offset by +1 from the common HIV-2 ROD^2^sequence after residue 8. We purified the protein to homogeneity (Figure S1A-S1E). Incubation of the purified protein with small unilamellar vesicles (SUVs) and IP6 resulted in the formation of CLPs marked by increased turbidity and the ability to pellet the CLPs via centrifugation. Imaging of the CLPs by negative stain EM revealed liposomes decorated with a repeating lattice similar to what had been observed with HIV-1 CLPs^44^(Figure S2A). Pelleted CLPs could be resuspended in a range of buffer pH values (6.0-8.5) and of salt concentrations (0 to 1 M NaCl) and maintain assembly, though with decreasing efficiency at lower or higher pH or high salt (Figure S2B-E). HIV-2 CA assembled poorly with only dNTPs or without any polyanions, similar to HIV-1 (Figure S2F and S2G). CA assemblies were stable for several days on ice or at 4°C, the limits of which have not been determined.

### High-resolution lattice structures of templated HIV-2 CLP assemblies by cryo-EM

Liposome-templated HIV-2 CLP assemblies were well-behaved as cryo-EM samples (Figure 1A), allowing for the determination of high-resolution structures (Figure 1 and Table S1). A cryo-EM map of the mature-like HIV-2 CA hexamer with C6 symmetry was resolved to 3.25 Å resolution (Figure 1B and S3). In initial stages of map classification and reconstruction, a subset of pentamers were observed adjacent to the central hexamer (Figure S3). Pentamer containing particles were re-processed with C5 symmetry to reconstruct an HIV-2 CA pentamer map to 2.97 Å resolution (Figure 1C and S3). In the final stages of processing the pentamer maps, a small portion of pentamer-containing particles were identified in assemblies with the appearance of T=1 pentamer icosahedra^38^(Figure 1A). These particles were re-picked and processed using icosahedral symmetry, yielding a 1.98 Å resolution map of the HIV-2 pentamer icosahedra (Figure 1D, 1E, and S4). Consistent with the high resolution, excellent density is observed and we can confidently build an atomic model of the CA molecule in the assembly (Figure 1F).

**Figure 1:**
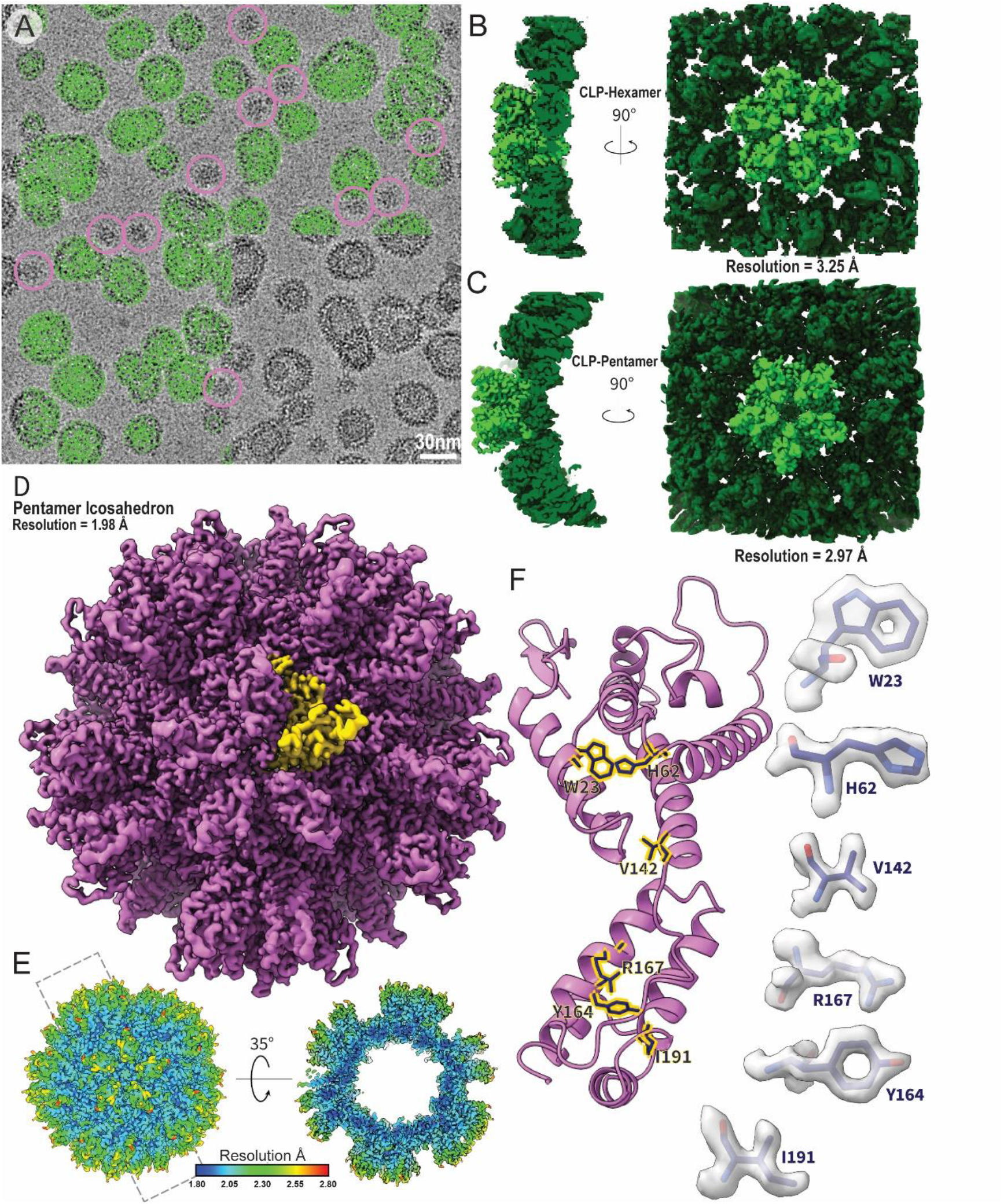
Assembly and structural analysis of liposome-templated HIV-2 CLPs **A**. Example cryo-EM micrograph of liposome-templated HIV-2 CLPs. Green circles mark example particles picked for 3D reconstruction. Pink circles mark example particles picked for icosahedral assemblies. **B**. Cryo-EM map of the HIV-2 CA hexamer. **C**. Cryo-EM map of the HIV-2 CA pentamer. **D**. Cryo-EM map of the micelle-templated HIV-2 pentamer icosahedron. Yellow highlights a single CA monomer. **E**. Local resolution map of the micelle-templated HIV-2 pentamer icosahedron. **F**. Cartoon representation of a CA monomer from the pentamer icosahedron. Sticks correspond to selected residues with corresponding map density highlighted to the sides.

Globally, both hexamer and pentamer maps were similar to those of HIV-1 CA assemblies. ^30,38,39,44^We observed density for IP6 in the central pores of both HIV-2 CA hexamers and pentamers, with clear density for R18 sidechains (Figure 2A and 2B). Density for a lower IP6 was also clear, though in the hexamer it was less well-resolved than the upper density and likewise, K25 sidechain density was more modest. As had been posited for the HIV-2 capsid lattice, the central pore appeared most similar to the open N-terminal β-hairpin conformation of the HIV-1 CA hexamer central pore,^18^despite that HIV-1 CA occupies the closed conformation in this pH range^54^ (Figure S5A). For both pentamers and hexamers, one of the most striking features of the map when compared with HIV-1 was stronger density of the Cyclophilin A (CypA)-binding loop (Figure 2C and S5B).

**Figure 2:**
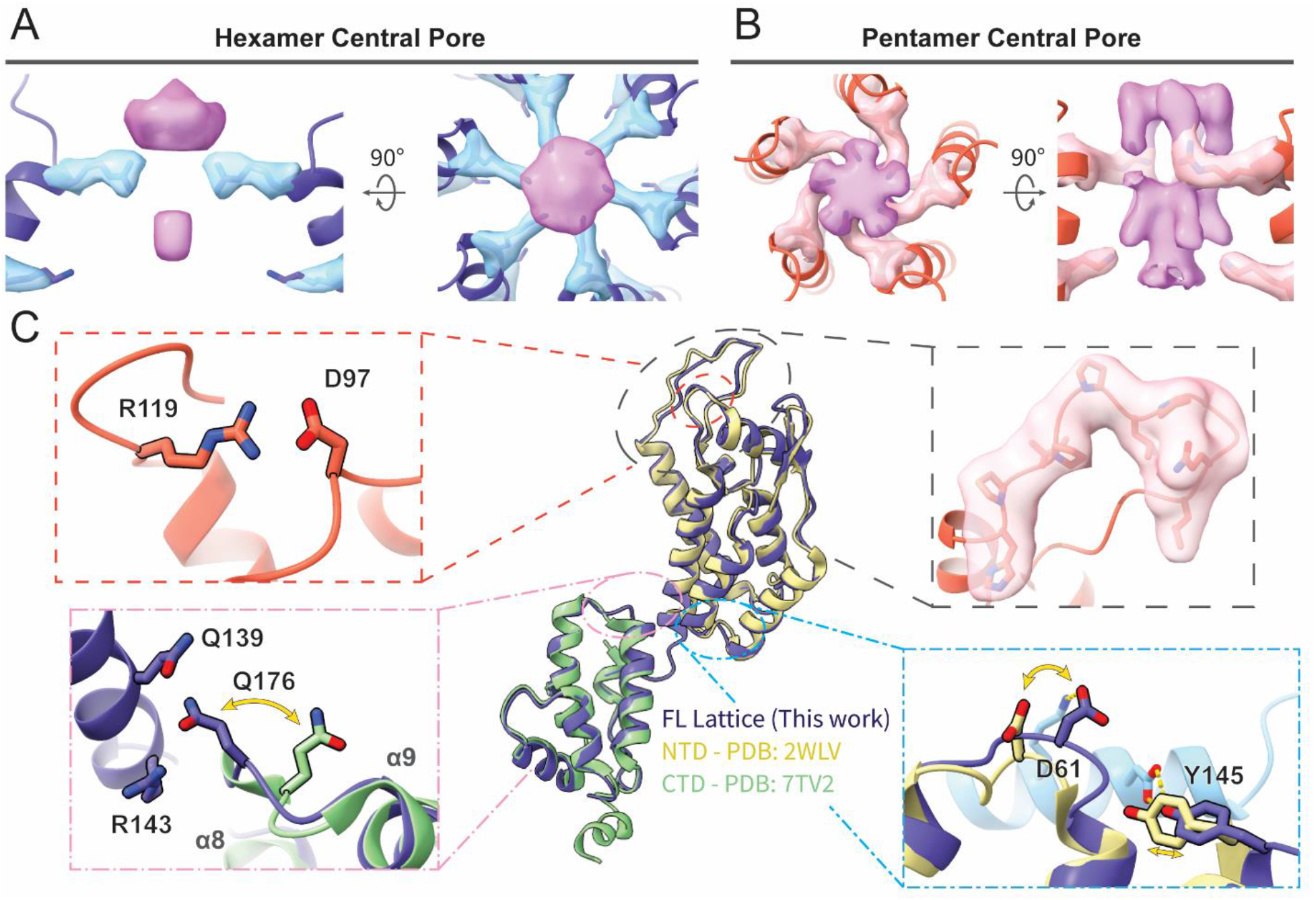
Atomic models of the mature HIV-2 CA lattice assembly. **A, B**. Top and side views of the IP6 density in the central pore of the HIV-2 CA hexamer (A) or pentamer (B), along with density for residues R18 and K25 (stick). The helices lining the pore are shown in cartoon representation. **C**. Cartoon representation of the CA protomer in the hexamer (blue) aligned with previous crystal structures of HIV-2 CA NTD^40^ (beige) and CTD ^41^ (green). Insets highlight structurally conserved salt bridge between GL-AN D97 and R119 (upper left), cryo-EM density of the CypA-binding loop (upper right), swinging out of the _176_QTD_178_ loop of the CTD upon ordered engagement with NTD in the CA lattice (lower left), and shift of the _58_CVGDH_62_ loop of the NTD and adjacent residues to accommodate adjacent protomer contact (light blue; lower right).

### The HIV-2 CA lattice exhibits distinctions from corresponding HIV-1 structures

Atomic models of HIV-2 CA could be confidently built into all maps and included all residues except those C-terminal to helix 11 (Table S2 and Figure S6A). The hexamer NTD aligned closely with the previously determined crystal structure of HIV-2 CA NTD^40^(∼0.9 Å RMSD; Figure 2C). This included structural conservation at the rare E97D mutation of GL-AN which retains formation of a salt bridge with R119 (Figure 2C, upper left), hypothesized to stabilize the CypA-binding loop.^18^Differences between the two structures largely occurred at protomer interfaces, either with adjacent chains or with the CTD (Figure 2C). Likewise, the hexamer CTD and previous HIV-2 CA CTD crystal structure^41^aligned closely (∼0.9 Å RMSD) (Figure 2C). However, the loop between helices 8 and 9 is shifted in the hexamer structure to allow Q176 in the CTD to contact residues Q139 and R143 in the NTD (Figure 2C, lower left). Furthermore, the previously described position of helix 12 in the CTD would clash with adjacent chains in our structure (Figure S6B-S6E)

The HIV-2 models reported here aligned closely with previously determined HIV-1 structures, both when comparing hexamers^34^ (∼1.7 Å RMSD) and pentamers^39^ (∼1.1 ÅRMSD; Figure S7A). As such, HIV-2 CA protomers also adopt significant conformational differences to accommodate forming both the pentamer and hexamer. However, considerable details underlying this structural plasticity differ. These differences also compound into identifiable changes in CA assemblies with the central pore diameter expanding by ∼2.7 Å in the hexamer^34^ and by ∼1.6 Å in the pentamer^39^ compared to HIV-1 at the Cα position of the R18 ring (Figure 3A). Comparing regions of the capsomers, the arrangement of NTD cores of HIV-2 is generally wider. In both viruses, CA NTD-NTD inter-protomer contacts are mediated by the N-terminal three-helix bundle and most sidechain interactions are conserved or similar. ^52^ However, while in HIV-1 the shift to the pentameric conformation excludes helix 3 from inter-chain contacts,^38,39^ helix 3 of HIV-2 CA remains engaged at the assembly interface with helix 2 from the adjacent chain through residues well-conserved within HIV-2, but divergent compared to HIV-1 (Figure 3B and S7C). Specifically, we observed an exchange of polar contacts as the hydrogen bonding potential of helix 2 residues Q41 and E45 are met either by Y50 and Q54 in the hexameric conformation or N57 and Q54 in the pentamer, allowing a change in register as helix 3 slides along helix 2.

**Figure 3:**
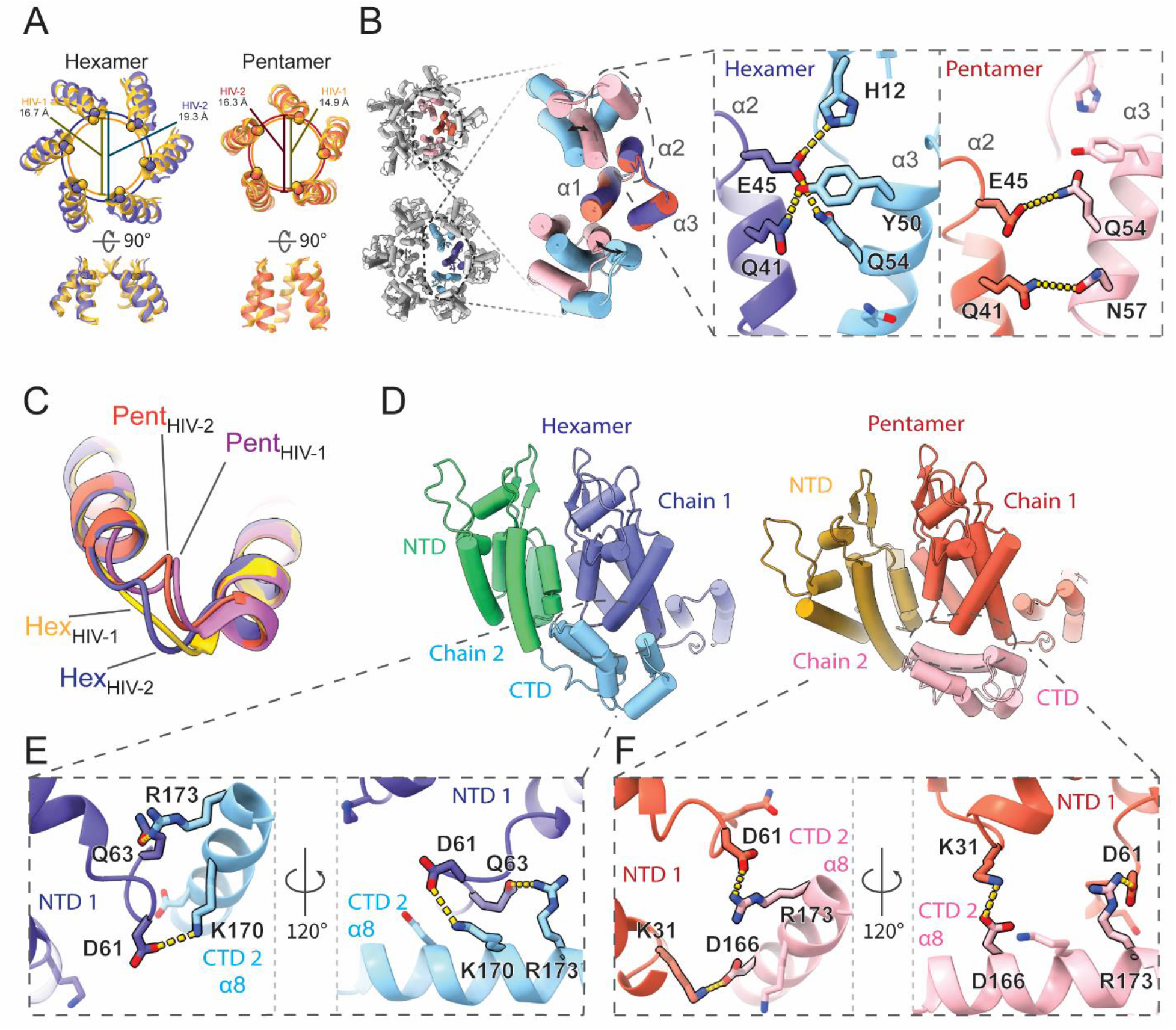
Unique intra-hexamer/pentamer contacts in the HIV-2 CA lattice. **A**. Alignment of the central pore of HIV-2 models with HIV-1 hexamer (PDB: 4XFX)^34^or HIV-1 pentamer (PDB: 8CKW)^39^. The overlapping circles delineate pore sizes as marked by R18 Cα distances. **B**. NTD-NTD intra-oligomer contacts are mediated by the N-terminal 3-helix bundle. Insets highlight exchange of hydrogen bonds between helices 2 and 3 to allow sliding of helix 3 between hexameric and pentameric conformations. **C**. Comparison of the _58_TVGG_61_ (HIV-1) and _58_CVGD_61_ (HIV-2) loops. While there is a shift between hexamer and pentamer forms in HIV-2, the loop structures stay similar and comparable to the HIV-1 pentamer _58_TVGG_61_ loop conformation. HIV-1 atomic models from PDBs 4XFX^34^(hexamer) and 8CKW^39^ (pentamer). **D**. The NTD-CTD contact area also shifts to accommodate the hexamer/pentamer transition. **E, F**. Sidechain details demonstrating the unique transition of polar contacts between HIV-2 CA NTD (chain 1) residues K31, D61, Q63 and residues in helix 8 of the adjacent CTD (chain 2).

While the _58_TVGG_61_ loop of HIV-1 has been described as a molecular switch between pentamer and hexamer,^38,39^ the _58_CVGD_61_ loop of HIV-2 does not demonstrate such a significant structural shift, though it is still structurally distinct between the two conformations (Figure 3C). The _58_CVGD_61_ loop of both conformations in HIV-2 is more similar to the pentameric conformation of HIV-1,^39^ though no 3_10_ helix is formed. Instead, D61 is engaged in distinct contacts with the CTD from the adjacent chain. Similar to the alternating NTD contacts, this interface permits a shift in helical register to accommodate either the hexameric or pentameric form (Figure 3D). In the hexamer, D61 and Q63 of one CA NTD form interactions with residues of the adjacent CA CTD, K170 and R173 (Figure 3E and S7D). On shifting to the pentamer, D61 replaces Q63 to ionically interact with adjacent CTD residue R173, while K31 of the NTD contacts the adjacent D166 (Figure 3F and S7E). These contacts are unique compared to HIV-1.^34,39^ Furthermore, we do not observe notable remodeling of the hydrophobic core of the NTD between hexamer and pentamer and the M39G mutation in HIV-2 precludes the ability of that residue to contribute to shifting knob-in-hole packing against adjacent chain helices as had been described for HIV-1. ^38^ In addition, we did not observe that residue M66 exhibits significant structural alteration between the oligomer states, different from what had been described as a gating mechanism for hexamer-pentamer transition in HIV-1 ^38,39^ (Figure S7F).

### Atomic details of FG-peptides binding to HIV-2 CA assemblies

With reproducible assemblies of HIV-2 CA amenable to structural study, we sought to resolve binding of FG-peptides derived from Nup153 and CPSF6 to identify differences in interactions compared to HIV-1 capsid.^39,55-60^ Additionally, since the conformational change of residue M66 contributes to exclude the binding of FG peptides to HIV-1 CA pentamers, we were curious whether the FG-peptides could bind to the HIV-2 CA pentamer that does not have steric occlusion by M66 rearrangement.

Binding of the Nup153 peptide (residues 1411-1425/1464-1475)^60^ was detected by co-sedimentation assays pelleting the liposome-templated CA assembly (Figure S8A). CA-SUV assemblies mixed with either peptide remained well behaved under both negative stain EM and cryo-EM imaging conditions (Figure S8B). In addition, induced assembly of apparent HIV-2 CA nanotubes with the introduction of Nup153 peptide was observed (Figure S8B). Following cryo-EM analysis, high-resolution structures of the HIV-2 CA hexamer (2.98 Å resolution) and pentamer (2.99 Å resolution) in the presence of the Nup153 peptide and of the hexamer (3.16 Å resolution) and pentamer (2.82 Å resolution) in the presence of the CPSF6 peptide (residues 313-327) ^58^ were determined (Figure 4 and S9).

**Figure 4:**
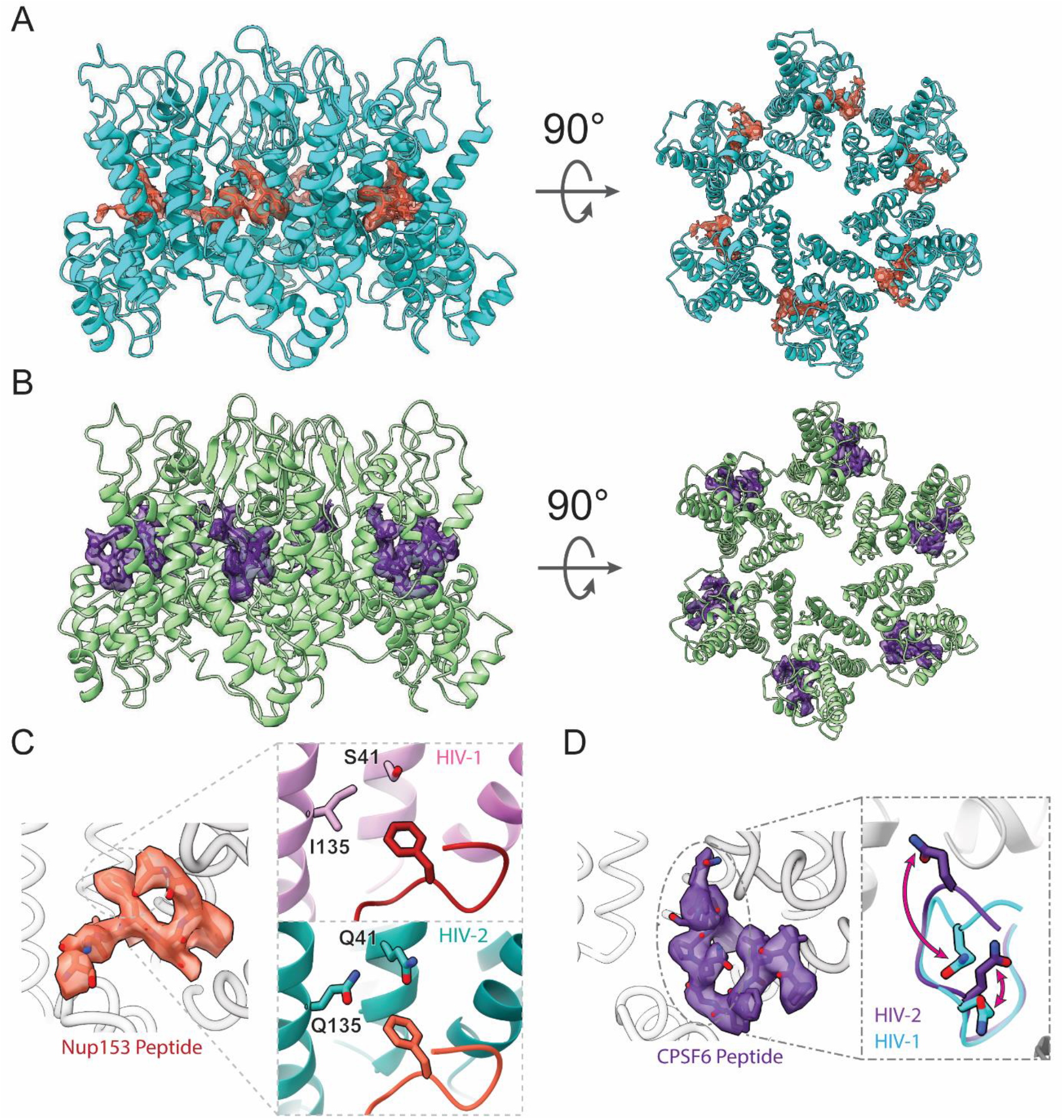
Binding of FG-peptides to HIV-2 CA hexamers. **A**. Cryo-EM reconstruction of the Nup153 FG-peptide binding to the HIV-2 CA hexamer. Nup153 density and model are in orange and HIV-2 CA in cyan. **B**. Cryo-EM reconstruction of the CPSF6 FG-peptide binding to the HIV-2 CA hexamer. CPSF6 density and model are in indigo and HIV-2 CA in green. **C**. Details of the atomic model built into the Nup153 FG-peptide density. The inset shows the N-terminal F of the FxFG peptide of Nup153 binds to a less hydrophobic pocket in HIV-2 CA hexamer with conserved polar residues, compared to those in HIV-1 (PDB: 8CKY). ^39^ **D**. Detailed atomic model built into the CPSF6 peptide density. The inset shows well-resolved structure for two glutamine residues interacting with backbone amide groups of HIV-2 CA, which have different conformations in the CPSF6 peptide/HIV-1 CA structures (PDB: 7SNQ).

Nup153 FG-peptide density was readily observed in the FG pockets of the HIV-2 CA hexamers (Figure 4A), but not of pentamers (Figure S10B). The density was strongest inside the FG pocket and extended to contact the CTD of the adjacent CA chain. An atomic model of 8 residues of the Nup153 peptide could be confidently built (Figure 4C; Table S2). Consistent with the high conservation of FG pocket residues between HIV-1 and -2, the model of Nup153 peptide in HIV-2 aligned closely with the known structure of Nup153 peptide in HIV-1 CA hexamer ^39^ (∼0.5 Å RMSD). This is true in spite of some well-conserved divergences between the two viruses along the adjacent CA NTD, such as a more polar ^*58*^ character of the HIV-2 CA FG pocket, which may have otherwise been expected to interact poorly with the first Phe of the Nup153 FxFG peptide (Figure 4C). Consistently, the PRODIGY web server ^61-63^ predicted similar binding energies for the Nup153 peptide to both CA assemblies, with -7.2 kcal/mol to HIV-1 CA and -7.4 kcal/mol to HIV-2 CA.

Similarly, CPSF6 FG-peptide density was readily observed in the FG pockets of HIV-2 CA hexamers (Figure 4B), but not pentamers (Figure S10C). This density was nearly as strong as main chain CA density and appeared to mainly contact one CA chain. An atomic model of 11 residues of the CPSF6 peptide could be confidently built (Figure 4D; Table S2). The FG segment of the peptide bound similarly to Nup153 peptide and was overall similar to the structure determined previously of CPSF6 peptide with HIV-1 CA hexamer ^58^ (∼0.4 Å RMSD). Interestingly, we observed clear interaction between two Gln residues of the CPSF6 peptide with backbone amide groups of the HIV-2 CA. These residues are also available in HIV-1 CA with similar conformations, but their interactions with the CPSF6 peptide was not observed previously (Figure 4D). ^58^ Consequently, binding energy prediction by the PRODIGY web server ^61-63^ predicted tighter binding of the CPSF6 peptide to HIV-2 CA (-9.0 kcal/mol) compared to HIV-1 CA (-5.4 kcal/mol). Despite resolving high-resolution structures of pentamers in the same data sets containing FG peptide-bound hexamers, no density for either FG-peptide was observed binding to the HIV-2 CA pentamer, like HIV-1, even in the absence of steric hindrance caused by the M66 rearrangement.

## Discussion

The viral capsid is indispensable to lentiviral infection and defines various sites of distinguishing host interactions between HIV-1 and HIV-2. ^5,16-26^ Leveraging an approach that has produced mature CA lattice assembly in HIV-1, ^44^ we have determined structures of the HIV-2 CA lattice assembly at high resolution containing both hexameric and pentameric conformations. These structures shared significant similarity with previously determined crystal structures of individual HIV-2 CA domains ^40,41^ and with corresponding HIV-1 CA protomers, ^34,38,39,44^ but diverging at inter-domain or inter-protomer interfaces or at locations of genetic divergence with HIV-1. These assemblies offer an accurate and reproducible representation of the mature HIV-2 capsid lattice, providing an efficient tool to study interactions with host factors.

While globally similar to HIV-1, these structures reveal distinctive regions in the HIV-2 CA lattice. The expanded central pore does not appear to affect the coordination of IP6. However, it may have implications on IP6 binding properties at the R18 or K25 rings, and thus on the rate of dNTP diffusion into the enclosed capsid. Additionally, the ordering of the CypA-binding loop provokes questions about the mechanism underlying differential interaction with an array of host factors, including CypA and RANBP2/Nup358. ^23,40,64^ Further, we identified diverging mechanisms to stabilize hexamer or pentamer formation. HIV-2 CA assemblies appear to lack the rearrangement of the hydrophobic core reported of HIV-1 capsomers. ^38,39^ Indeed, the divergence of M39G in HIV-2 CA may preclude adopting the same mechanism as has been described for HIV-1. Instead, the HIV-2 CA structures exhibit several additional polar contacts between adjacent chains, particularly between NTD helices 2 and 3, which could explain how they modulate forming pentamers and hexamers. This subtle rearrangement suggests that the _58_CVGD_61_ loop of HIV-2 functions more gradually, rather than the switch mechanism proposed for HIV-1. ^38,39^ In accordance with this model, we observe icosahedral pentamer-only particles without kinetically trapping a molecular switch in a particular state by mutation. ^38^

With reliable atomic models of both HIV-1 and HIV-2 capsomers revealing details of complex formation, we can better analyze the evolutionary space of primate lentiviruses. Focusing on residues at the infaces of the assemblies, two trends quickly become apparent. First, M39 is well-conserved among primate lentiviruses, with the M39G mutation appearing only in the monophyletic grouping of closely related HIV-2 and SIV_smm_. This group also exhibits mutations V/A26L and S/T41Q nearby in space (Figure 5B). L26 appears to partially compensate for conversion of M39 to glycine in maintaining the hydrophobic interface and would otherwise clash with M39 if both were present (Figure 5C). This is consistent with assembly and maturation issues exhibited by HIV-2 G39M revertant mutants. ^65^ Q41 permits hydrogen bonding with residues on helix 3. A hydrogen bonding partner of Q41, Q54, appears in a slightly broader monophyletic grouping, adding in the SIV_rcm_ and SIV_tan_ branch. It is possible that Q54 arose first and was followed by Q41 to interact and stabilize helix 2-3 interactions, permitting the M39G and V26L mutations.

**Figure 5:**
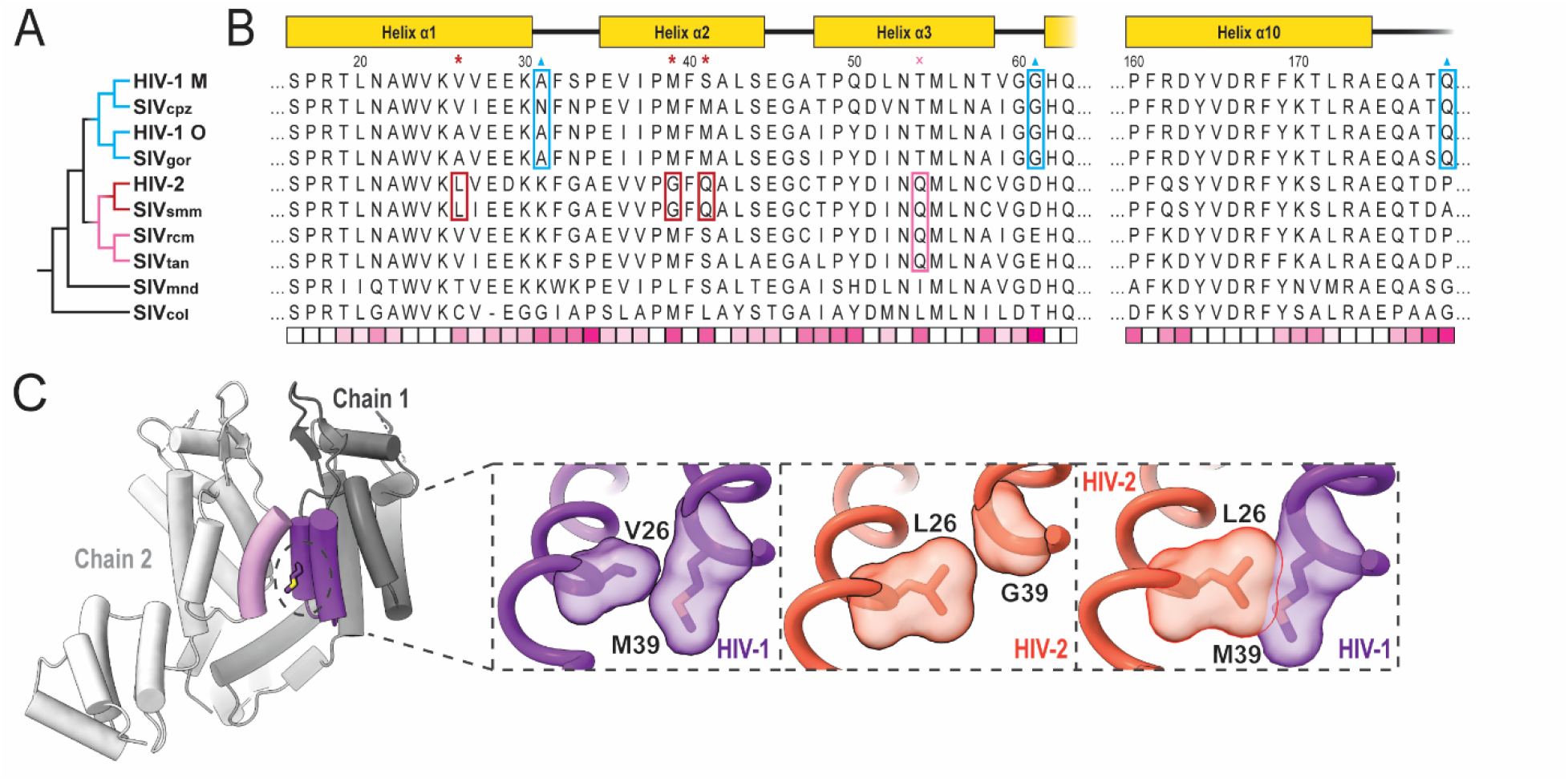
Mutations implicated in pentamer/hexamer formation among primate lentiviruses. **A**. Phylogenetic tree of selected primate lentiviruses based on CA amino acid sequence similarity. **B**. Multiple sequence alignment of primate lentivirus CA amino acid sequences. Sequences are either from a prior defined group consensus ^66^ or consensus derived from sequences deposited at the Los Alamos National Laboratory database. ^67^ Schematic of HIV-1/HIV-2 domain architecture at the top. Residues marked by red asterisks and boxes are specific to the HIV-2/SIV_smm_ group. Residue marked by pink cross and box are specific to a broader group of HIV-2-related lentiviruses. Residues marked by blue triangles and boxes are specific changes among the HIV-1-related group. The bottom row displays relative conservation per position based on BLOSUM-80 scoring. ^68^ Highly conserved positions appear whiter while more divergent positions appear more pink. **C**. Example of the change at position 39 in HIV-1 ^34^ and HIV-2. Left and central insets highlight the native context of positions 26 and 39 in HIV-1 and HIV-2 with surface representations. Right inset shows aligned HIV-1 and HIV-2 structures with clash between L26 and M39 if without additional changes.

Sequence elements of the pentamer switch loop _58_TVGG_61_ of HIV-1 CA appear uniquely in the HIV-1/SIV_cpz_/SIV_gor_ monophyletic group. The HIV-2-related group as well as more ancestrally related SIV_mnd_ exhibit well-conserved D/E61 which we identified forming ionic interactions with K170/R173 in HIV-2 assemblies. It is unclear how this potential interaction would affect the switch remodeling of the _58_TVGG_61_ loop, but D/E61G is particular to the HIV-1-related group (Figure 5B). Further, this HIV-1 related grouping also appears to lose K31 as a potential CTD contact in the pentamer, but gains P/A179Q to hydrogen bond with Q63 in the adjacent chain, potentially as a trade-off to accommodate other structural changes. Taken altogether, these series of mutations provide clues for different scenarios regarding the evolution of the mechanisms of achieving CA pleomorphism.

With high resolution HIV-2 CA lattice structures, we can identify 14 surface residues with conserved change of chemical character between HIV-1 and HIV-2 which lack clear structural reason for divergence (Figure S11; Table S3). This may imply elements of this set potentially contribute to differential host factor binding. Despite these and other differences, the mode of FG-peptide binding for those we tested appeared largely conserved. Furthermore, despite lack of M66 gating in HIV-2, we did not observe density for either CPSF6 or Nup153 peptide in the FG pocket of CA pentamers, indicating another mechanism is responsible for reduced binding to HIV-2. Future studies, leveraging the efficient in vitro assembly system and functional experiments, will answer more questions regarding HIV-2/HIV-1 capsid lattice structure formation and host factor binding, presenting many new opportunities to better understand the biology and evolution of these viruses.

## Acknowledgements

This work was funded by the National Institutes of Health through grants T32GM008283, U54AI170791, and P50AI150481. Cryo-EM data was collected with the support of staff at the Brookhaven National Laboratory – Laboratory for BioMolecular Structure (LBMS) and the Yale CryoEM Resource. LBMS is supported by the DOE Office of Biological and Environmental Research (KP1607011). Computational resources used are maintained by the Yale Center for Research Computing. We thank the Xiong lab members for discussions and support of this work, particularly Dr. M. Meuser, F. Arizaga, and S. Balaji. We also thank Dr. R. Dick for technical advice and suggestions.

## Author Contributions

M.C., C.F., and Y.X. conceived this project and designed the study; C.F. prepared the functionalized liposomes, expressed and purified the HIV-1 protein, assembled SUV-templated HIV-1 CLPs and prepared EM samples, collected cryo-EM data, and performed cryo-EM reconstruction; M.C. expressed and purified the HIV-2 protein; M.C. assembled CLPs and prepared samples with discussion from C.F.; M.C. and F.L. performed assembly and cosedimentation assays; M.C. performed negative stain EM imaging; C.F. and C.W. performed cryo-EM data collection; C.F. and X.Y. performed cryo-EM reconstruction; M.C. performed model building with discussion from C.W., C.F., and Y.X.; M.C., C.F., and Y.X. analyzed the data; M.C., C.F., and F.L. wrote the paper; M.C. and C.F. created the visualizations; M.C., C.F., and Y.X. edited the paper. All of the authors discussed the results and commented on the manuscript.

## Declaration of Interests

The authors declare no competing interests.

## STAR Methods

### Lead Contact and Materials Availability

The plasmid pET11a GL-AN HIV-2 CA-GSSHHHHHH was produced for this study and is available upon request. Further information and requests for the plasmid, resources, and reagents should be directed to and will be fulfilled by the Lead Contact, Dr. Yong Xiong.

### Protein Expression and Purification

A hexahistidine tag with Gly-Ser-Ser linker was cloned onto the C-terminus of GL-AN HIV-2 CA in pET11a vector. GL-AN HIV-2 CA-GSS-6xHis was expressed in BL21(DE3) competent cells. Cells were grown to OD_600_ ∼0.8 in Terrific broth before induction with 500 μM isopropyl β-d-1 thiogalactopyranoside at 18°C for 16 h and then collected by centrifugation. Cells were resuspended in lysis buffer (50 mM Tris·HCl, pH 8.0 at 4°C; 500 mM NaCl; 0.2 mM tris(2-carboxyethyl)phosphine (TCEP); 1 x Halt™ protease inhibitor cocktail) and lysed in a microfluidizer. The lysate was clarified via centrifugation before bulk protein was precipitated from the supernatant with 35% ammonium sulfate (stirring at 4°C for 1 h). Precipitated protein was collected by centrifugation and then resuspended in NiNTA Buffer A (50 mM Tris·HCl, pH 8.0; 350 mM NaCl; 20 mM imidazole; 0.2 mM TCEP). The resuspended protein was clarified by centrifugation and then loaded onto a 5mL hand-packed column of NiNTA agarose resin (Qiagen). Protein was eluted with NiNTA Buffer B (50 mM Tris·HCl, pH 8.0; 350 mM NaCl; 300 mM imidazole; 0.2 mM TCEP) in a stepwise fashion. The eluted peak was then dialyzed against cation exchange Buffer A (25 mM Na·HEPES, pH 6.9; 50 mM NaCl; 0.2 mM TCEP) overnight (stirring at 4°C). The dialyzed protein was clarified by centrifugation and loaded onto a 5 mL HiTrap SP HP column (Cytiva) for cation exchange chromatography. The protein began eluting around 25% of the way through a 50 mM to 1 M NaCl gradient. Protein was concentrated to 1 mM concentration and snap frozen with final buffer containing 300 mM NaCl.

### Liposome Preparation

The preparation scheme was adapted form Highland et al., 2023. ^44^ Chloroform stock solutions of 1,2-dioleoyl-sn-glycero-3-[(N-(5-amino-1-carboxypentyl)iminodiacetic acid)succinyl] nickel salt (DGS-NiNTA) and 1,2-dioleoyl-sn-glycero-3-phosphocholine (DOPC) were purchased from Avanti Polar Lipids. Cholesterol was purchased from Thermo Scientific Chemicals and resuspended at 5mg/mL in chloroform. Lipids were mixed at an 85:10:5 DOPC:DGS-NiTNA:Cholesterol ratio using a volumetric flask so that the final composition of the lipid mixture is roughly 95% DOPC, 2%DGS-NTA, and 3% cholesterol. The lipid mixture was transferred to a round bottom flask and dried by rotary evaporation for 5 hours. The resulting “lipid cake” was resuspended in aqueous buffer (25mM HEPES pH 7.4, 150mM KCl) to a final concentration of 13 mM by gentle agitation on a rotovap. The resuspended “lipid cake” was allowed to hydrate overnight at room temperature. Large unilamellar vesicles (LUVs) were prepared by extruding the hydrated lipid cake through a 100 nm polycarbonate membrane 100 times at 37°C. Small unilamellar vesicles (SUVs) were prepared by sonication with a nanoprobe tip at 50% amplitude for 15 minutes of processing time (10 second pulse, 50 seconds off). Lipid preparations were verified by negative stain EM before use in assembly assays (Figure S12).

### CA Assembly by Liposome Templating and Cosedimentation

Capsid like particles (CLPs) were assembled by mixing the following components to the given final concentrations: 2.5 mM IP6, 3.7 mM SUV or LUV lipids, and 250 μM GL-AN HIV-2 CA-GSS-6xHis. The particles were assembled during a 15 min incubation at 37°C and allowed to recover for 5 min at room temperature. To study the binding of Nup153 peptide or CPSF6 peptide and HIV-2 CLPs, the FG-peptide was added to the already-assembled CLPs following recovery. The FG-peptide·CLP complex was then allowed to form during a 30 min incubation at room temperature.

To test binding via cosedimentation, samples were pelleted down by centrifugation at 16k rcf for 20 m using a room temperature tabletop microcentrifuge. The supernatant was then drawn off, which was typically around the original sample volume. The pellet was washed with an equivalent volume of desired buffer before centrifuging again at 16k rcf for 5 m. Wash buffer was removed and the pellet was resuspended in equivalent buffer volume for subsequent analysis by SDS-PAGE or EM imaging.

### Negative Stain EM of Liposomes and CLPs

3.5 µL of liposome sample was applied to a glow-discharged negative stain EM grid (EMS, carbon on 300-mesh copper) for 1 min. The grid was washed 1X with buffer (25mM HEPES pH 7.4, 150 mM KCl) then stained in 2% uranyl acetate for 1 min and blotted with filter paper. For CLPs, a 3.5 μL sample diluted 1:4 in buffer was deposited onto glow-discharged (25 mA for 30 s) 400 mesh carbon-coated copper grids (Electron Microscopy Services) for 1 min before blotting. Grids were stained with 2% uranyl acetate solution (Electron Microscopy Services) for 90 s before blotting. Imaging was conducted on a 120 kV Talos™ L120C TEM with CETA CMOS camera.

### Cryo-EM Sample Preparation

A volume of 3.5 μL of CLPs was deposited onto glow-discharged (15 mA, 45 s) Quantifoil™ R 2/1 200 mesh copper grids. Sample was dual-side blotted for 5.5 – 7.5 s by a Vitrobot (ThermoFisher) with chamber at 100% humidity and then plunge-frozen in liquid ethane.

### Cryo-EM Data Collection and Processing

Cryo-EM data were collected at Brookhaven National Laboratory (BNL) and Yale University cryo-EM facilities. In each case, movies were collected using a 300 kV Titan Krios (ThermoFisher) equipped with an energy filter and a K3 direct detector (Gatan). Movies were collected using EPU (ThermoFisher) (BNL) or SerialEM ^69^ (Yale) at a physical pixel size of 1.068 Å (Yale) or 1.07 Å (BNL) in super-resolution mode with a total dose of 50 electrons per Å ^2^ and a target defocus of -0.8 to -2.0 µm.

Image processing was performed using CryoSPARC. ^70-72^ For the HIV-2-SUV dataset, 6,037 movies were subjected to patch motion correction (bin2, pixel size 1.068 Å) and CTF estimation. Initial manual picking of 1,213 particles followed by 2D classification yielded usable templates for template-based particle picking. Separate template picking jobs were performed to pick “top” and “side” views of the capsid lattice. After manual inspection, duplicate particles were removed and the remaining 13,209,858 particles were extracted using a box size of 104 pixels (bin4, pixel size 4.272 Å). Extracted particles were subjected to several rounds of 2D classification and selection. The remaining 3,065,617 particles were subjected to C1 homogenous refinement with EMD-3465 ^35^ lowpass filtered to 40 Å. Subsequent heterogenous refinement of the 3,065,617 particles using 4 identical reference volumes (EMD-3465^35^lowpass filtered to 40 Å yielded two hepta-hexamer classes (total 1,572,733 particles), one class containing pentamer (1,030,843 particles), and a junk class.

Pentamer containing particles were used for a C5 ab-initio reconstruction which resulted in a map that features a central pentamer adjacent to five hexamers. At this point, particles were re-extracted in box size of 416 pixels (pixel size 1.068 Å) without binning. These particles were used for homogenous refinement, 3D classification, non-uniform refinement, and several rounds of iterative local refinement paired with per-particle CTF refinements (global and local CTF estimation), ultimately yielding a 2.97 Å map (346,785 particles).

Likewise, hexamer containing particles were used for a C6 ab-initio reconstruction, which resulted in a map that features a central hexamer surrounded by six others. At this point, particles were re-extracted in box size 416 (pixel size 1.068 Å) without binning. These particles were used for homogenous refinement, 3D classification, non-uniform refinement, ^73^ and several rounds of iterative local refinement paired with per-particle CTF refinements (global and local CTF estimation), ultimately yielding a 3.25 Å resolution map (603,473 particles).

For the HIV-2-Nup153 peptide dataset, 3,732 movies were collected and motion correction, CTF estimation, and initial particle picking were performed as previously described. After template picking, 11,931,864 particles were subjected to multiple rounds of heterogenous refinement using 3 identical volumes (EMD-3465 ^35^ lowpass filtered to 40 Å) and 2 junk volumes (EMD-3465 lowpass filtered to 100 Å). After five rounds of heterogenous refinement the total percentage of particles sorted to junk dropped below 1% and the refinement was considered to have converged. Pentamer-containing particles were aligned to the apo HIV-2 pentamer volume using homogenous refinement and re-extracted in box size 180 (pixel size 1.068 Å). These particles were used for homogenous refinement, non-uniform refinement, and several rounds of iterative local refinement paired with per-particle CTF refinements (global and local CTF estimation), ultimately yielding a 2.98 Å resolution map (1,259,201 particles). A parallel procedure for the hexamer-containing particles, aligning to the apo HIV-2 hexamer volume, yielded a 2.99 Å resolution map (1,494,376 particles).

For the HIV-2-CPSF6 peptide dataset, 5,506 movies were collected and motion correction, CTF estimation, and initial particle picking were performed as previously described. After template picking, 23,033,474 particles were subjected to multiple rounds of heterogenous refinement as described above. Similarly, subsequent parallel processing of pentamer-containing and hexamer-containing particles yielded a 2.82 Å resolution pentamer map (1,906,465 particles) and a 3.16 Å resolution hexamer map (2,537,344 particles).

Pentamer icosahedrons were observed in all HIV-2-SUV samples at a lower abundance. As the HIV-2-CPSF6 sample had the highest particle concentration, we focused our analysis on this dataset. For the pentamer icosahedrons, manual picking of 314 particles yielded clear 2D classes, which were subsequently used for template picking. After manual inspection, 690,629 particle picks were extracted in a box of 102 pixels (bin 4, pixel size 4.272 Å) and subjected to several rounds of 2D classification to remove non-icosahedral lattice particles. 75,595 final selected particles were subjected to ab-initio reconstruction with icosahedral symmetry imposed. The resulting maps featured a micelle coated in 12 pentamers.

Subsequently, the particles were re-extracted in box 384 (pixel size 1.068 Å) with no binning and subjected to homogenous refinement, non-uniform refinement,^73^and several rounds of iterative local refinement paired with per-particle CTF refinement (global and local CTF estimation), ultimately yielding a 2.18 Å resolution map (74,821 particles). At this point, the particles were subjected to reference-based motion correction^71,74^(super-resolution pixel size 0.712 Å) followed by non-uniform refinement with CTF fitting (tetrafoil, spherical aberration, anisotropic mag),^73^defocus estimation and positive curvature EWS correction yielding a 1.98 Å resolution map (74,821 particles).

### Atomic Model Building and Refinement

An initial model was prepared in Coot^75,76^ by docking the cryo-EM structure of pentameric WT HIV-1 CA from IP6-stabilized CLPs (PDB: 8CKW)^39^ into the pentamer-centered cryo-EM map of HIV-2 CA templated on SUV. Coot was then used to manually rebuild the structure to match the density. Iterative rounds of Phenix^77^ Real Space Refinements and manual adjustments in Coot were then performed until model quality was evaluated as acceptable. This model was then used as a reference in the icosahedral pentamer cryo-EM map and rebuilt and refined as above. As the icosahedral pentamer model was derived from a higher resolution map, it was then used as initial reference for all subsequent models, including hexamers after adjusting the relative orientation between CA NTD and CTD, which were also refined as above. Model quality was evaluated by the Phenix ^77^Comprehensive validate (cryo-EM) suite including MolProbity. ^78^RMSD measurements for structure comparisons were performed using the MatchMaker tool in ChimeraX. ^79-81^

## Data Availability

Cryo-EM density maps have been deposited in the Electron Microscopy Data Bank with the following accession codes: EMD- (HIV-2 CA hexamer), EMD- (HIV-2 CA pentamer), EMD- (HIV-2 CA icosahedron), EMD- (HIV-2 CA hexamer with Nup153 peptide), EMD- (HIV-2 CA pentamer from Nup153 peptide dataset), EMD- (HIV-2 CA hexamer with CPSF6 peptide), and EMD- (HIV-2 CA pentamer from CPSF6 peptide dataset). Atomic coordinates have been deposited in the Protein Data Bank with the following accession codes: (HIV-2 CA hexamer), (HIV-2 CA pentamer), (HIV-2 CA icosahedron), (HIV-2 CA hexamer with Nup153 peptide), and (HIV-2 CA hexamer with CPSF6 peptide).

The plasmid for GL-AN HIV-2 CA-GSS-6xHis is available upon request. Any additional information required for reanalysis is also available upon request.

## Notes

### Competing Interest Statement

The authors have declared no competing interest.

