## Supplemental Information for "Structural insights into HIV-2 CA lattice formation and FG-pocket binding revealed by single particle cryo-EM"

**Supplemental Figures and Tables**

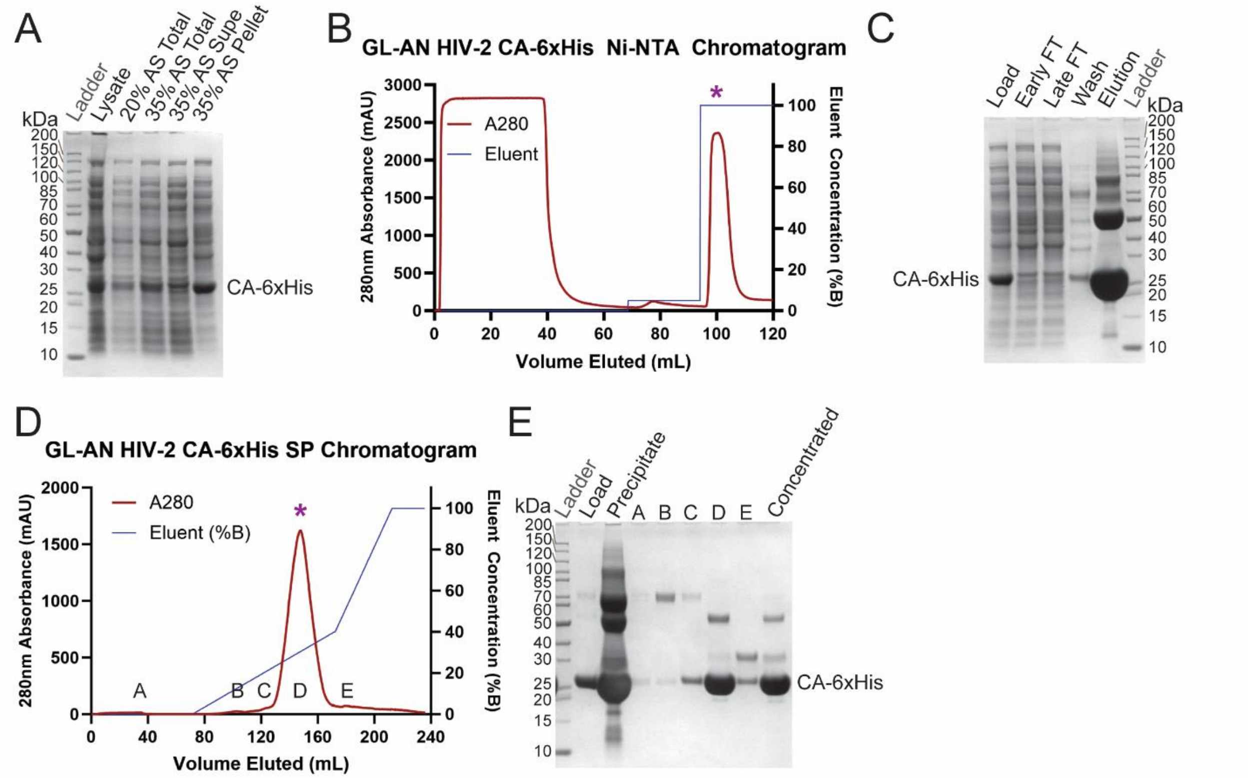


*Figure S1: Preparation of GL-AN HIV-2 CA-6xHis and Liposomes.*

***A***. *SDS-PAGE result of GL-AN HIV-2 CA-6xHis (27 kDa) expression lysate and ammonium sulfate (AS) precipitation steps.* ***B***. *Ni-NTA chromatogram of resuspended 35% AS pellet of GL-AN HIV-2 CA-6xHis.* ***C***. *SDS-PAGE result of Ni-NTA chromatography.* ***D***. *Cation exchange chromatogram from HP SP column of dialyzed Ni-NTA peak fraction (purple asterisk in B) sample of HIV-2 CA-6xHis.* ***E***. *SDS-PAGE result of cation exchange chromatography (purple asterisk in C) and protein concentration.*

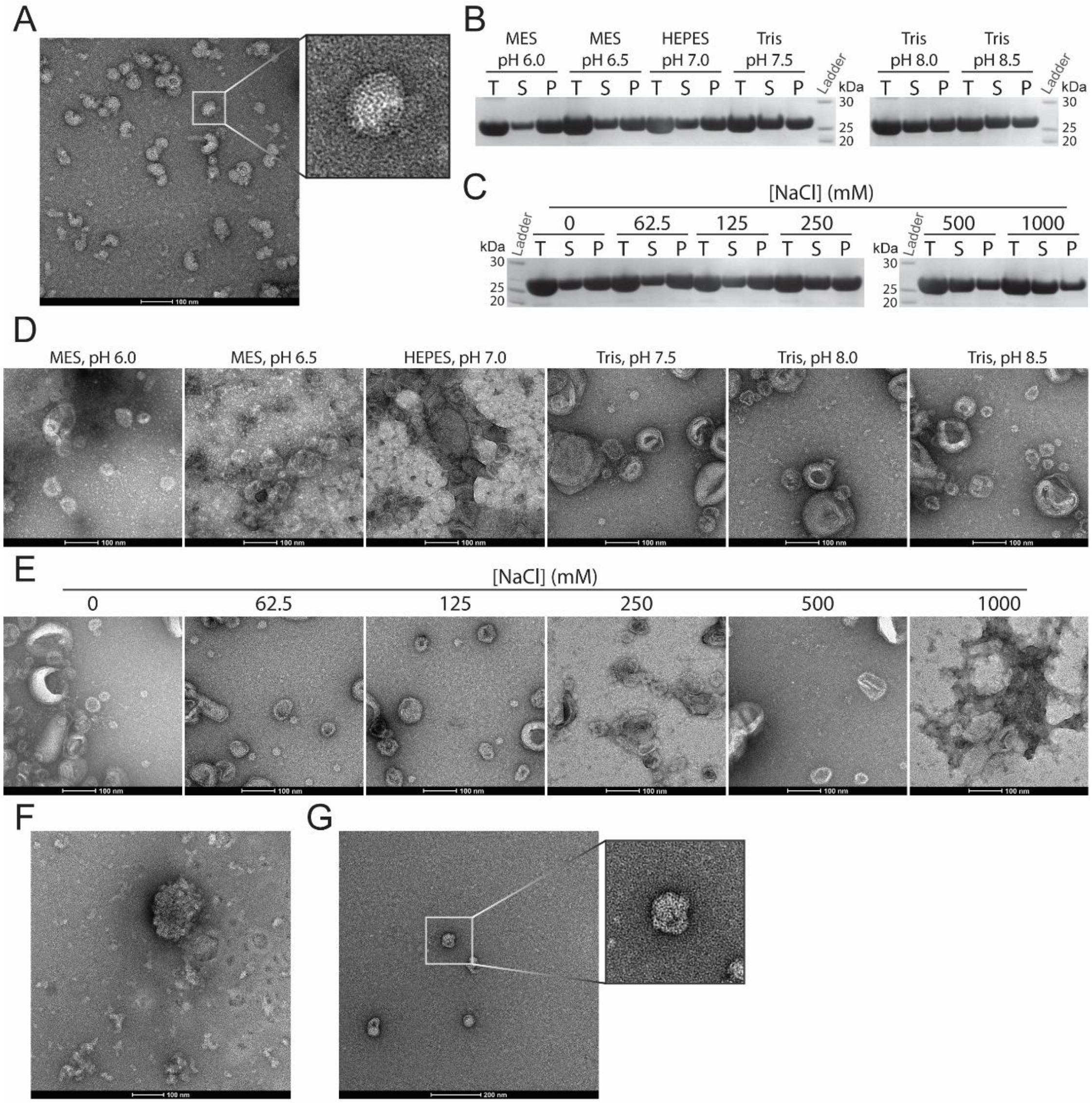


*Figure S2: Assembly of HIV-2 capsid-like particles by liposome templating.*

***A**. Representative negative stain (NS) EM micrograph of SUV-templated CLP assemblies at 73,000 x magnification. **B**. SDS-PAGE gels of sedimented LUV-templated HIV-2 CLPs following resuspension in buffers of varying pH, 150 mM NaCl. T lanes represent Total CA; S, Supernatant CA; P, Pelleted CA. **C**. SDS-PAGE gels of sedimented LUV-templated HIV-2 CLPs following resuspension in buffers of varying NaCl concentration, 50 mM Tris, pH 7.5. Lane labeling as B. **D**. Representative NS EM micrographs at 73,000 x magnification of resuspended pellet samples from B. **E**. Representative NS EM micrograph at 73,000 x magnification of resuspended pellet samples from C. **F**. Representative NS EM micrograph at 73,000 x magnification of SUV-templated CLP assembly without any polyanions - no well assembled CLPs were observed. **G**. Representative NS EM micrograph at 52,000 x magnification of SUV-templated CLP assembled in the presence of dNTPs instead of IP6. Infrequent assemblies were observed.*

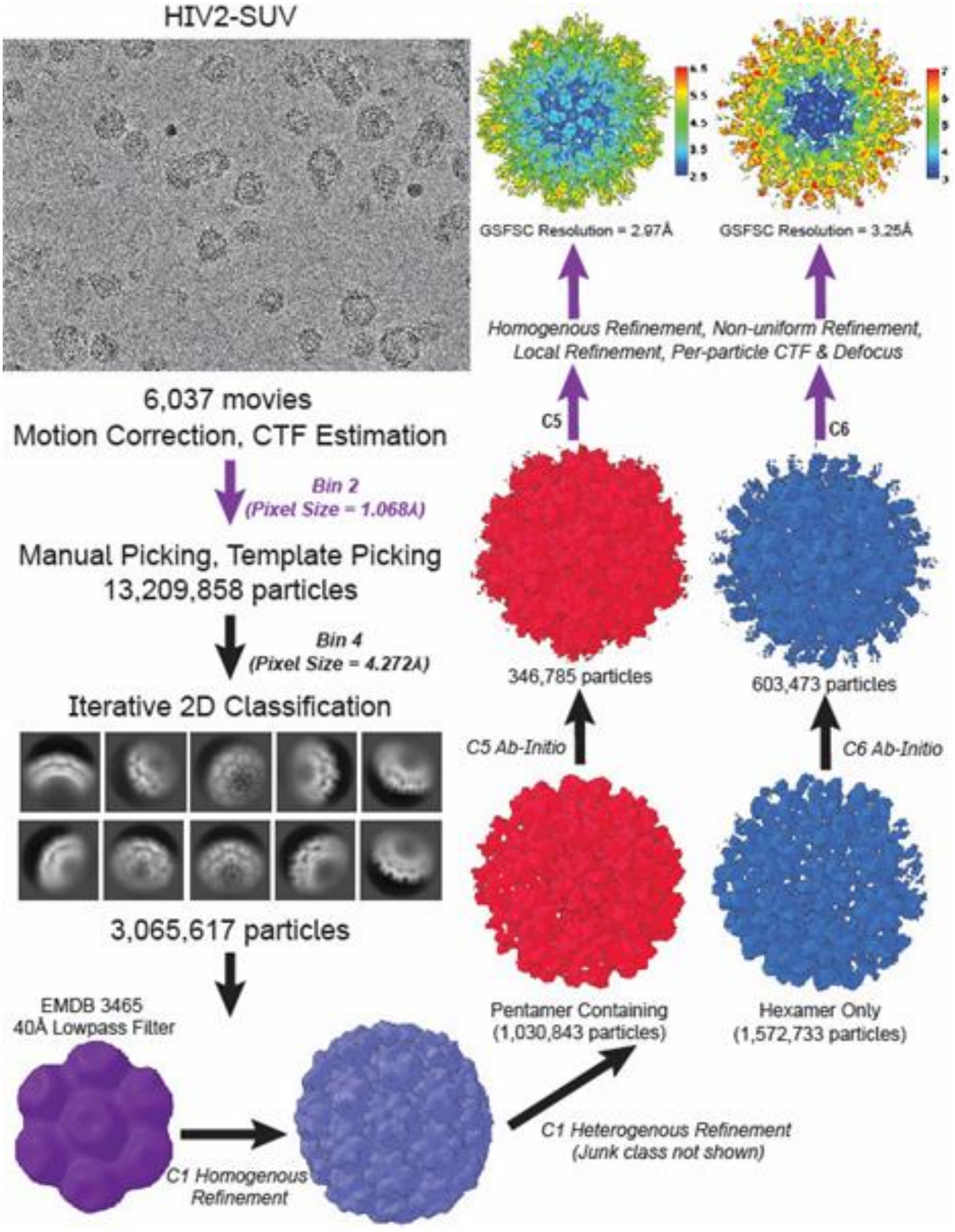


*Figure S3: Cryo-EM data processing flowchart of HIV-2 CA hexamer and pentamer.*

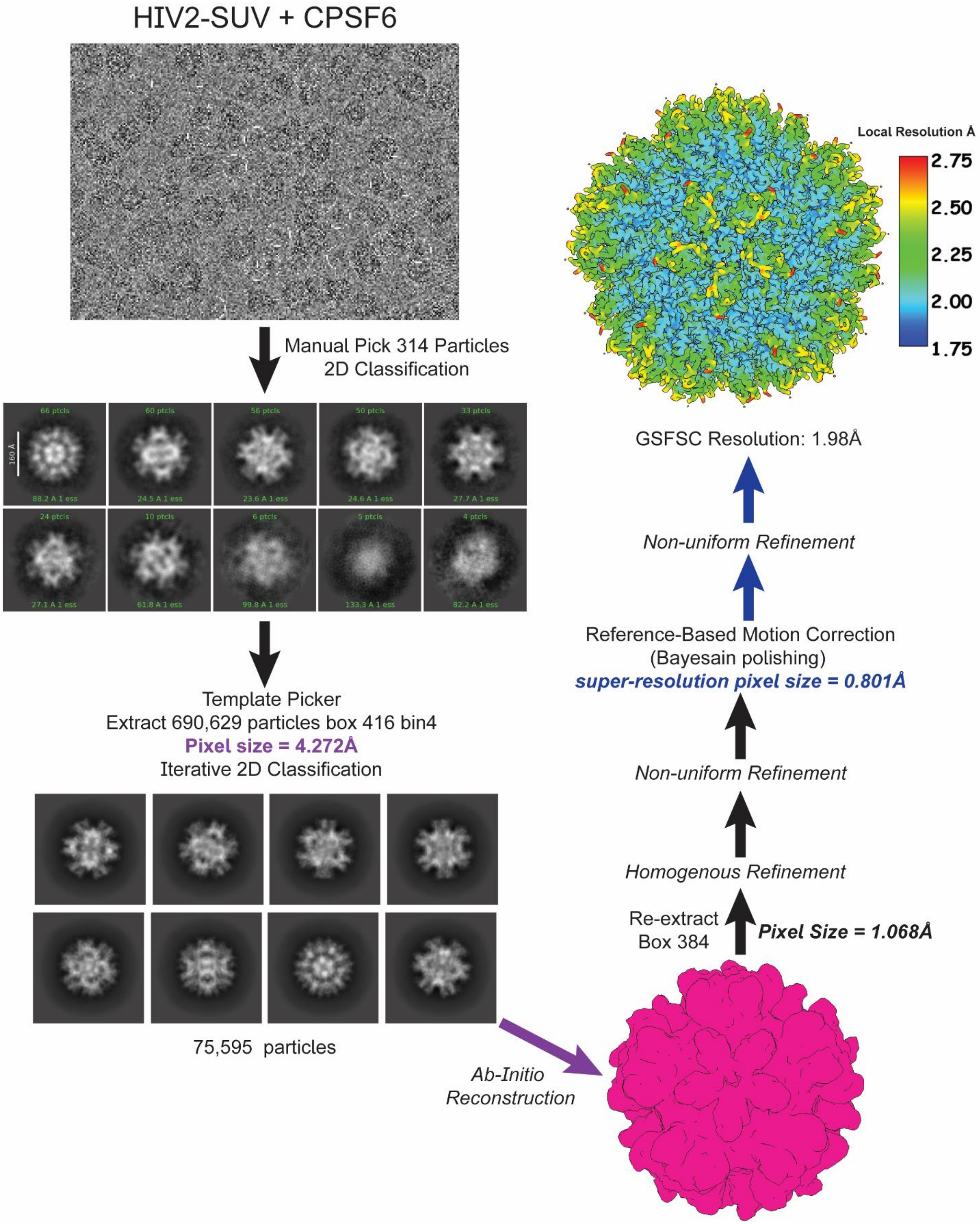


*Figure S4: Cryo-EM data processing flowchart of HIV-2 CA icosahedra.*

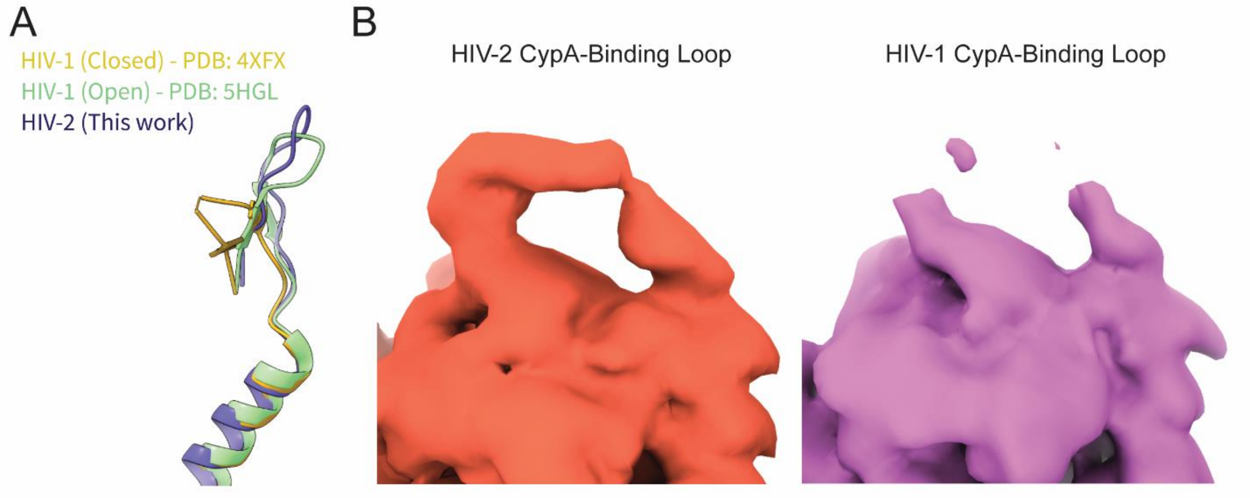


*Figure S5: Comparison of CA loop regions between HIV-2 and HIV-1.*

***A***. *Comparing the N-terminal β-hairpin loop of HIV-2 CA with those of closed*[34] *and open*[54] *conformations of HIV-1 CA.* ***B***. *Comparison of cryo-EM density of the CypA-binding loop for HIV-2 and HIV-1 CA pentamer maps at similar contour levels.*

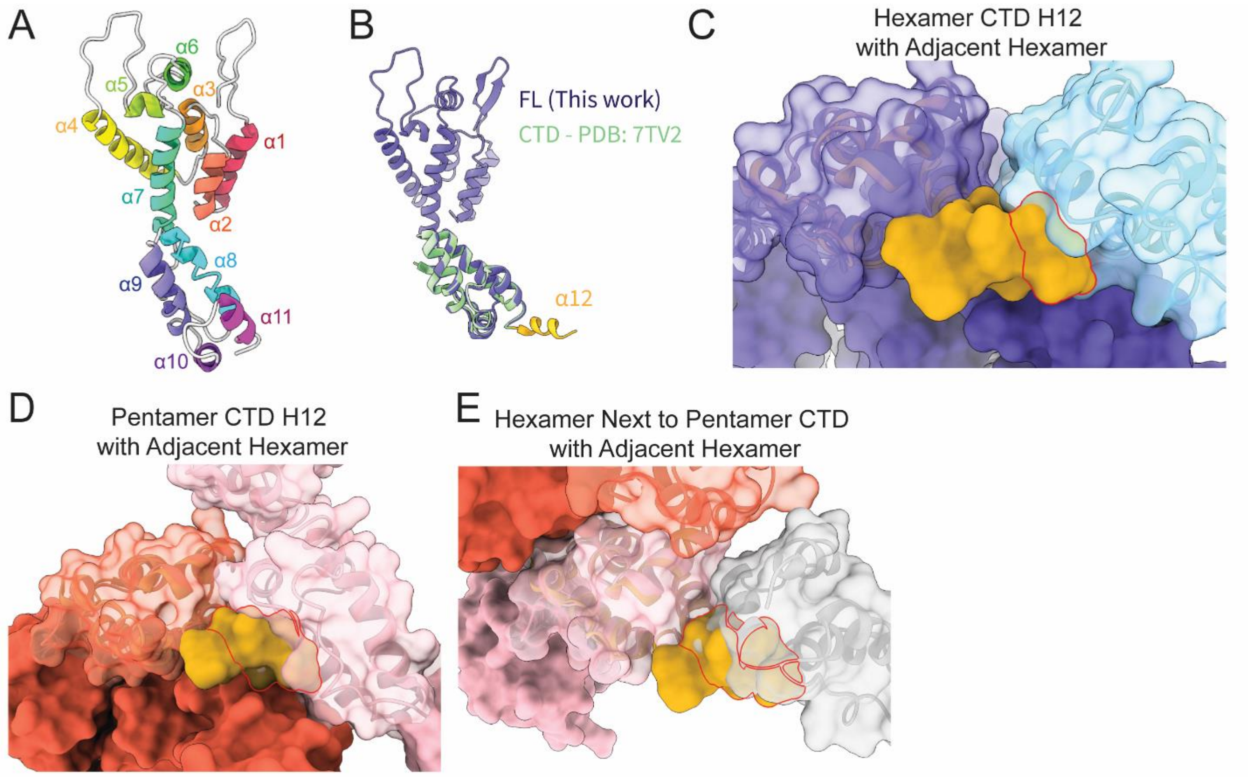


*Figure S6: Helix enumeration and potential clash of monomeric CA helix 12 in the lattice.*

***A**. Enumerating the α helices identified in HIV-2 CA, matching that for HIV-1 CA. **B**. Model alignment of FL HIV-2 CA (this work) with HIV-2 CA CTD with emphasis on helix 12 from the CTD structure.[41] **C**. Surface representation of HIV-2 CA CTD structure (7TV2)[41] aligned to the FL CA in the hexameric lattice. Aligned CA protomer in dark blue. Adjacent CA protomer in light blue. Helix 12 of the CTD structure in gold. Residues outlined in red clash with the adjacent chain. **D**. Surface representation of HIV-2 CA CTD structure (7TV2)[41] aligned to the FL CA in the pentameric lattice. Aligned CA protomer in red. Adjacent CA protomer in pink. Helix 12 of the CTD structure in gold. Residues outlined in red clash with the adjacent chain. **E**. Surface representation of HIV-2 CA CTD structure (7TV2)[41] aligned to the FL CA in a hexamer adjacent to a pentamer. Aligned CA protomer in red. Adjacent CA protomer in pink. Helix 12 of the CTD structure in gold. Residues outlined in red clash with the adjacent chain.*

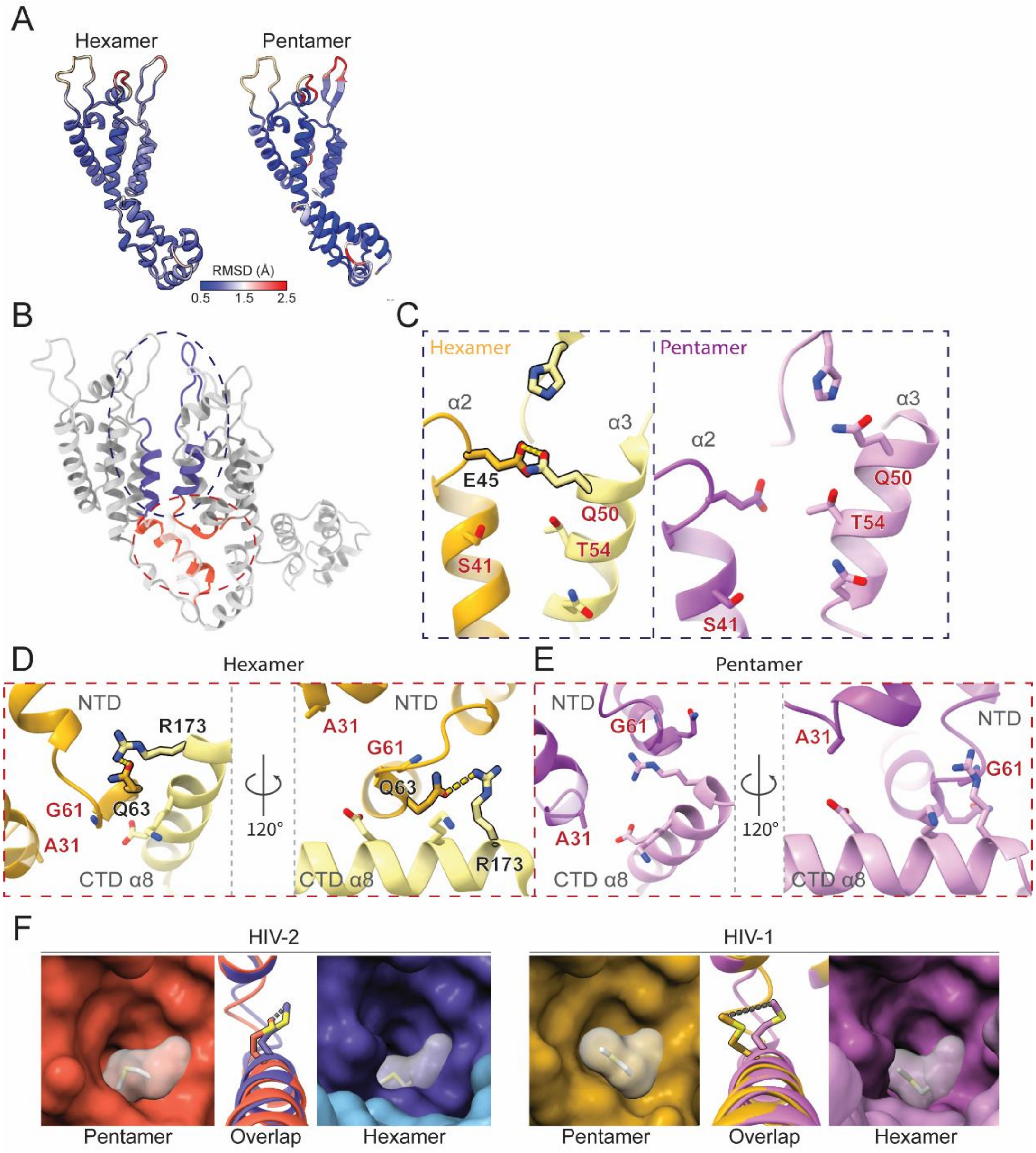


*Figure S7: Comparisons of contacts in HIV-2 and HIV-1 CA.*

***A***. *Alignment of HIV-2 hexamer and pentamer chains with corresponding ones of HIV-1,[34,39] colored by RMSD.* ***B***. *Model of adjacent CA promoters to orient NTD-NTD (blue) and NTD-CTD (red) contacts.* ***C***. *NTD-NTD contacts involving helices 2 and 3 in HIV-1 CA, focusing on residues with ionic interactions in HIV-2 CA (hex: 4XFX[34]; pent: 8CKW).[39] Residues labeled in red are divergent between HIV-1 and HIV-2.* ***D, E***. *NTD-CTD contacts in HIV-1 CA, focusing on residues with ionic interactions in HIV-2 CA (hex: 4XFX[34]; pent: 8CKW).[39] Residues labeled in red are divergent between HIV-1 and HIV-2.* ***F***. *FG pocket comparison between HIV-2 and HIV-1.[34,39] Surface representations of respective models are shown. M66 is shown in stick with white transparent surface, which also covers the flexible residue R70 (HIV-2) or K70 (HIV-1) (not shown).*

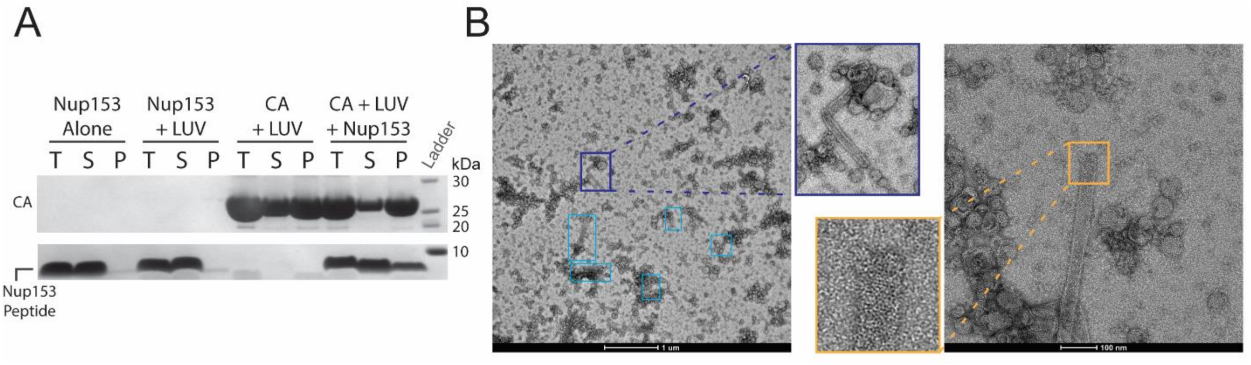


*Figure S8: Interaction of Nup153 peptide with HIV-2 CA.*

***A**. SDS-PAGE gel of HIV-2 CLP cosedimentation of Nup153 peptide. **B**. Negative stain EM micrographs demonstrating observed non-liposome-templated CA nanotubes assembled in the presence of Nup153 peptide. Left micrograph captured at 11,000 x magnification, blue boxes mark the positions of the nanotubes. Right micrograph at 73,000 x magnification revealing ordered lattice and lack of evidence of internal lipids in the nanotubes.*

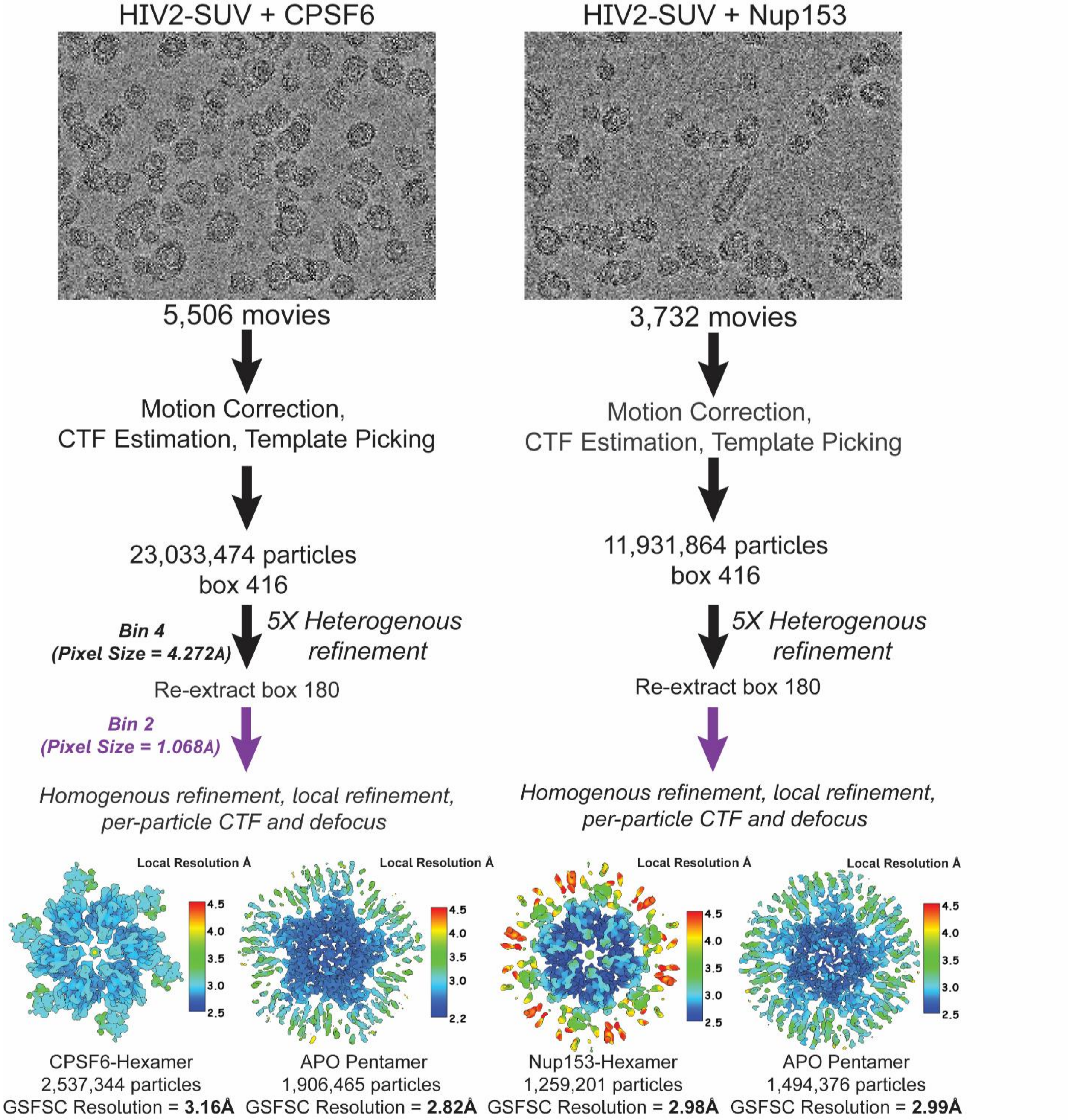


*Figure S9: Cryo-EM data processing flowcharts of HIV-2 CA hexamers and pentamers in the presence of either Nup153 peptide or CPSF6 peptide.*

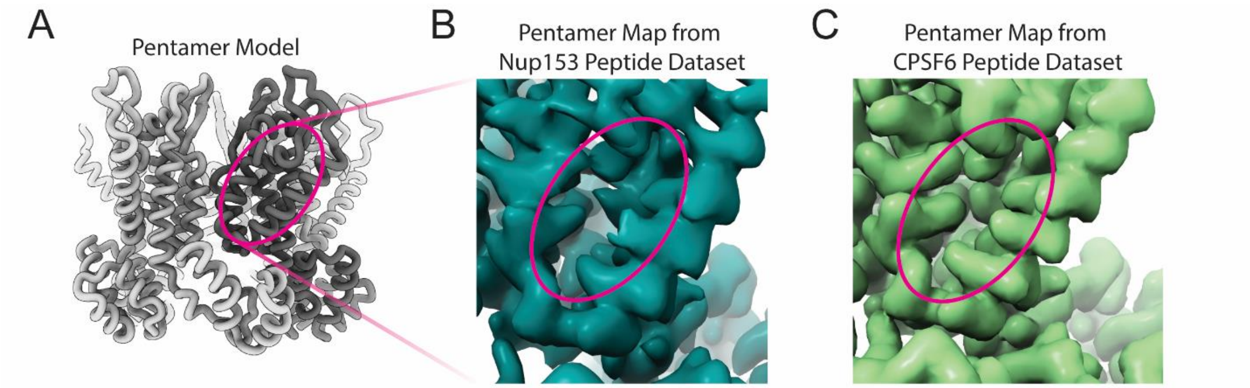


*Figure S10: Empty FG pockets in HIV-2 pentamers treated with FG peptides.*

***A****. HIV-2 CA pentamer model orienting the location of the FG pocket (magenta oval).* ***B****. Pentamer map from the same dataset deriving Nup153 peptide-bound hexamers revealing lack of density observed in the pentamer FG pocket.* ***C****. Pentamer map from the same dataset deriving CPSF6 peptide-bound hexamers revealing lack of density observed in the pentamer FG pocket.*

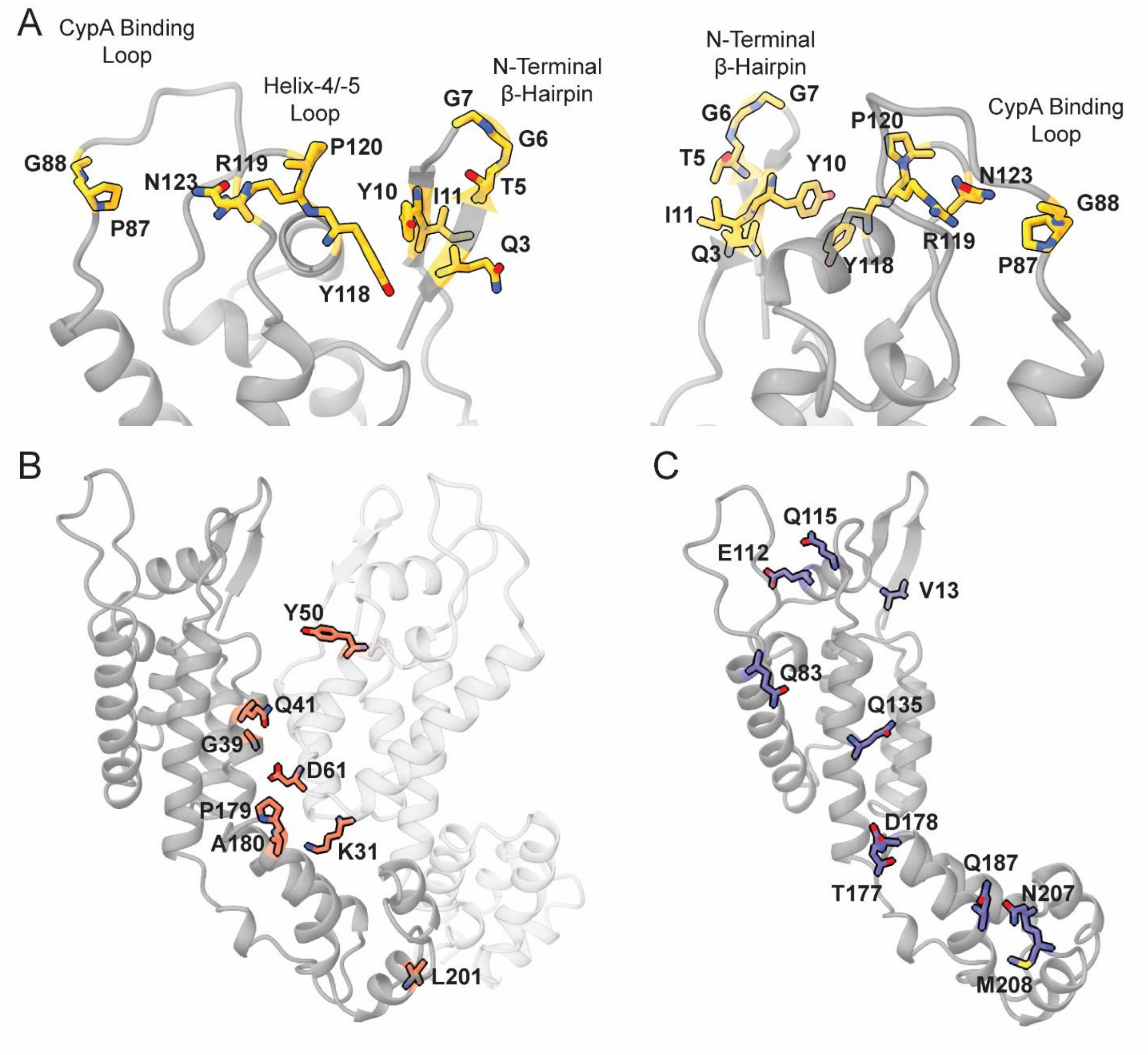


*Figure S11: Surface and interface residues of HIV-2 divergent from HIV-1.*

***A**. Divergent residues between HIV-2 and HIV-1 in CA N-terminal loop regions, defined as changes in chemical character or clear structural divergence. **B**. Divergent residues between HIV-2 and HIV-1 at inter-protomer interfaces. **C**. Divergent residues between HIV-2 and HIV-1 on the exterior mature capsid surface.*

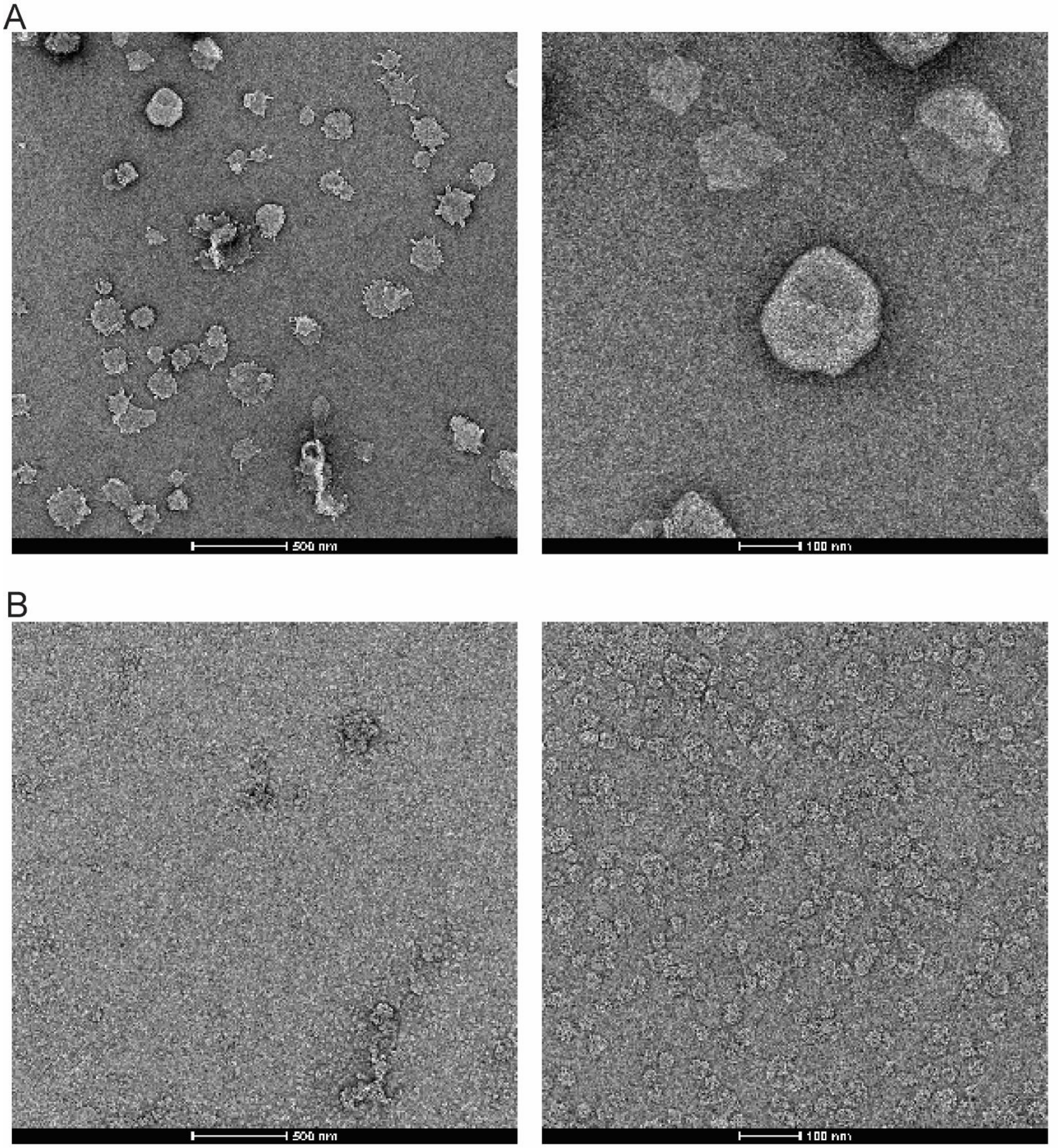


*Figure S12: Negative stain of liposome assemblies.*

***A**. Left: Low magnification image of large unilamellar vesicles after 100 extrusion passes. Right: High magnification image of a LUV. **B**. Left: Low magnification image of small unilamellar vesicles after 15 minutes of sonication processing time. Right: High magnification image of SUVs.*

| Particle | HIV-2 CA Hexamer | HIV-2 CA Pentamer | HIV-2 CA Icosahedron | HIV-2 CA Hexamer with Nup153 Peptide | HIV-2 CA Pentamer from Nup153 Peptide Data Set | HIV-2 CA Hexamer with CPSF6 Peptide | HIV-2 CA Pentamer from CPSF6 Peptide Data Set |
|---|---|---|---|---|---|---|---|
| **Microscope** | Titan Krios | Titan Krios | Titan Krios | Titan Krios | Titan Krios | Titan Krios | Titan Krios |
| **Detector** | K3 (Gatan) | K3 (Gatan) | K3 (Gatan) | K3 (Gatan) | K3 (Gatan) | K3 (Gatan) | K3 (Gatan) |
| **Magnification** | 81,000 x | 81,000 x | 81,000 x | 81,000 x | 81,000 x | 81,000 x | 81,000 x |
| **Voltage (keV)** | 300 | 300 | 300 | 300 | 300 | 300 | 300 |
| **Exposure ($e^-/Å^2$)** | 50 | 50 | 50 | 50 | 50 | 50 | 50 |
| **Super-resolution mode?** | Yes | Yes | Yes | Yes | Yes | Yes | Yes |
| **Acquisition Software** | SerialEM | SerialEM | SerialEM | SerialEM | SerialEM | SerialEM | SerialEM |
| **Number of movie frames** | 40 | 40 | 40 | 40 | 40 | 40 | 40 |
| **Defocus Range (µm)** | -0.8 to -2.0 | -0.8 to -2.0 | -0.8 to -2.0 | -0.8 to -2.0 | -0.8 to -2.0 | -0.8 to -2.0 | -0.8 to -2.0 |
| **Pixel Size (Å)** | 1.068 | 1.068 | 1.07 (super-resolution to 0.712) | 1.068 | 1.068 | 1.07 | 1.07 |
| **Symmetry Imposed** | C6 | C5 | I | C6 | C5 | C6 | C5 |
| **Final Particles** | 603,473 | 346,785 | 74,821 | 1,494,376 | 1,259,201 | 2,537,334 | 1,906,465 |
| **FSC Threshold** | 0.143 | 0.143 | 0.143 | 0.143 | 0.143 | 0.143 | 0.143 |
| **Resolution (Å)** | 3.25 | 2.97 | 1.98 | 2.98 | 2.99 | 3.16 | 2.82 |
| **Resolution Range (Å)** | 3.0 to 7.0 | 2.5 to 6.5 | 1.75 to 2.75 | 2.5 to 4.5 | 2.5 to 4.5 | 2.5 to 4.5 | 2.2 to 4.5 |
| **EMDB ID** | EMD-45758 | EMD-45759 | EMD-45676 | EMD-45760 | EMD-45762 | EMD-45761 | EMD-45763 |

*Table S1: Cryo-EM data processing statistics.*

| Model | HIV-2 CA Icosahedron | HIV-2 CA Hexamer | HIV-2 CA Pentamer | HIV-2 CA Hexamer with Nup153 Peptide | HIV-2 CA Hexamer with CPSF6 Peptide |
|---|---|---|---|---|---|
| **PDB ID** | 9CLJ | 9CNS | 9CNT | 9CNU | 9CNV |
| **Model Composition** | | | | | |
| **Chains** | 2 | 3 | 4 | 2 | 2 |
| **Non-hydrogen atoms** | 1,974 | 2,971 | 7,016 | 1,862 | 1,898 |
| **Protein residues** | 222 | 369 | 885 | 229 | 233 |
| **Water** | 130 | 0 | 0 | 0 | 0 |
| **IP6** | 2 | 2 | 2 | 2 | 2 |
| **Root mean squared deviations** | | | | | |
| **Bond lengths (Å)** | 0.012 | 0.013 | 0.012 | 0.013 | 0.013 |
| **Bond angles (°)** | 1.7 | 1.9 | 1.7 | 2.0 | 1.8 |
| **Validation** | | | | | |
| **MolProbity score** | 1.24 | 1.50 | 1.52 | 1.65 | 1.51 |
| **Clash score** | 1.1 | 4.9 | 4.5 | 6.9 | 4.3 |
| **Rotamer outliers (%)** | 1.6 | 0.6 | 0.7 | 0.0 | 0.0 |
| **Ramachandran plot validation** | | | | | |
| **Favored (%)** | 95.9 | 96.4 | 95.8 | 96.0 | 95.6 |
| **Allowed (%)** | 4.1 | 3.6 | 4.2 | 4.0 | 4.4 |
| **Outliers (%)** | 0.0 | 0.0 | 0.0 | 0.0 | 0.0 |
| ***Z*-score** | 1.2 | 2.7 | 1.2 | 2.2 | 1.6 |

*Table S2: Atomic model refinement and validation statistics.*

| Loops | | | Interfaces | | | Surface | | |
|---|---|---|---|---|---|---|---|---|
| **HIV-2** | **Position** | **HIV-1** | **HIV-2** | **Position** | **HIV-1** | **HIV-2** | **Position** | **HIV-1** |
| Q | 3 | V | K | 31 | A/G | V/I/M | 13 | Q |
| V/I | 5 | N | G | 39 | M | Q/A | 83 | L/V/T |
| G/A | 6 | L/V | Q | 41 | S/T | E/D | 112\|113 | Q |
| G | 7 | Q | Y | 50 | Q | Q/E | 115\|116 | G/A |
| Y | 10 | M | D | 61 | G | Q | 135 | I |
| T/S | 11 | V/I | P | 179 | Q | T | 177 | A |
| P | 87 | H/Q | A | 180 | E/D | D | 178 | T/S |
| _ | _ \| 88 | A | L | 201 | T/S | Q | 187 | E/D |
| F/Y | 118\|119 | T | - | - | - | M/I | 207 | P/T |
| R | 119\|120 | S/N/H | - | - | - | N | 208 | G/A |
| A/P | 120\| _ | _ | - | - | - | - | - | - |
| N | 122 | P | - | - | - | - | - | - |

*Table S3: Residues with divergent chemical character between HIV-2 and HIV-1.*

*Position number following canonical HIV-1 CA numbering and HIV-2 GL-AN CA numbering used elsewhere in this manuscript. Where numbering diverges, the left number describes HIV-2 location, the right describes HIV-1. Amino acids with > 10% prevalence across HIV-1 or HIV-2 populations are shown. An insertion/deletion not present in one chain is marked with an underscore ( _ ).*
